# MLL3 and MLL4 sustain hematopoietic stem cell multipotency by opposing a B-cell default state

**DOI:** 10.64898/2025.12.24.696404

**Authors:** Helen C. Wang, Ran Chen, Wei Yang, Rohini Muthukumar, Tingting Hu, Riddhi M. Patel, Emily B. Casey, Elisabeth Denby, Run Zhang, Guojia Xie, Kai Ge, Grant A. Challen, Jeffrey J. Bednarski, Jeffrey A. Magee

## Abstract

Hematopoietic stem cells (HSCs) and multipotent progenitors (MPPs) are sustained by networks of transcription factors and epigenetic regulators that prime lineage-specific programs yet maintain multipotency. Two epigenetic regulators, MLL3 and MLL4, play important but opposing roles in maintaining this balance. MLL3 promotes HSC differentiation, whereas MLL4 opposes differentiation. These opposing functions are essential for both normal homeostasis and leukemia suppression, yet it is not clear how MLL3 and MLL4 regulate HSC and MPP gene expression to control HSC/MPP fate decisions. To resolve these mechanisms, we performed an extensive series of single cell genomic studies after conditionally deleting *Mll3*, *Mll4* or both genes together. *Mll3* deletion had only limited effects on HSC/MPP enhancer networks at steady state, whereas *Mll4* deletion led to precocious activation of myeloid enhancers. Surprisingly, compound *Mll3/4* deletion eliminated all myeloid, erythroid and megakaryocytic potential within the hematopoietic hierarchy and caused all progenitors to rapidly default to a B-cell-like identity. These changes were accompanied by widespread inactivation of HSC/MPP enhancers and superenhancers, and ectopic activation of B-cell superenhancers. Disabling MLL3/4 histone methyltransferase activity did not recapitulate the pervasive changes in cell identity that were observed when MLL3 and MLL4 were fully inactivated, indicating that MLL3 and MLL4 activate HSC/MPP enhancers independently from their enzymatic activities. Our findings show that HSC/MPP multipotency requires sustained tension between MLL3/4-dependent enhancers that maintain myeloid, erythroid and megakaryocyte potential, and MLL3/4-independent enhancers that prime B-cell identity. MLL3 and MLL4 therefore serve as critical linchpins of multilineage hematopoiesis.

**KEY POINTS:**

- MLL3 and MLL4 act redundantly in HSCs to sustain transcription factor and enhancer networks that support multipotency
- Simultaneous loss of MLL3 and MLL4 drives hematopoietic progenitors into a uniform B-cell-like default state

## Introduction

Throughout life, blood cells arise from hematopoietic stem cells (HSCs) and multipotent progenitors (MPPs) that either self-renew or differentiate into myeloid, lymphoid, megakaryocytic, or erythroid progenitors.^1–6^ Maintaining this extensive degree of multipotency requires gene regulatory networks that prime several different lineage programs within a given stem cell while simultaneously preventing any one program from fully asserting itself, up until the time of lineage commitment. Gene regulatory programs that prime and maintain the multipotent state have been heavily scrutinized, and many critical transcription factors have been identified.^7–9^ For example, the transcription factor MECOM has recently been shown to prevent HSC differentiation by directly repressing genes that encode myeloid transcription factors, such as CEBPA.^10,11^ Likewise, combinatorial interactions between several transcription factors, including TAL1, LYL1, LMO2, GATA2, RUNX1, ERG and FLI1, serve to maintain HSCs.^9^ These transcription factors also regulate lineage-specific target genes at later stages of differentiation.^8^ Epigenetic regulators often interact with transcription factors to enforce cell fate decisions by covalently modifying histones or DNA, or by altering chromatin accessibility. These regulators play essential roles in maintaining normal HSC homeostasis and suppressing leukemic transformation, yet we have only a limited understanding of how they are deployed to coordinate HSC/MPP fate decisions.

MLL3 (also called KMT2C) and MLL4 (also called KMT2D) are two highly homologous epigenetic regulators that have established roles in balancing HSC self-renewal and differentiation. They therefore offer potential insights into how individual epigenetic regulators coordinate gene expression to either reinforce the multipotent state or promote differentiation.^12–14^ Each protein nucleates large chromatin-bound complexes that belong to the <u>Com</u>plex of <u>P</u>roteins <u>As</u>sociated with <u>S</u>ET1 (COMPASS) family (Figure 1A). COMPASS complexes bind enhancer elements to promote gene expression via histone modifications, transcriptional condensate formation and RNA polymerase pause-release.^15–20^ These activities drive cell identity changes across a range of tissues in all metazoan species, and they are essential for tumor suppression.^17,21–24^ However, in the hematopoietic system, MLL3 and MLL4 perform distinct, antagonistic functions that seem incongruent with their structural similarities and overlapping functions in other tissues. MLL3 restricts HSC self-renewal by promoting differentiation.^12,14^ In contrast, MLL4 sustains HSC self-renewal capacity by opposing differentiation.^13^ It is not clear how such structurally similar protein complexes antagonize one another, and the prior studies raise the question of whether MLL3 and MLL4 also perform overlapping, non-antagonistic functions in HSCs.

**Figure 1.**
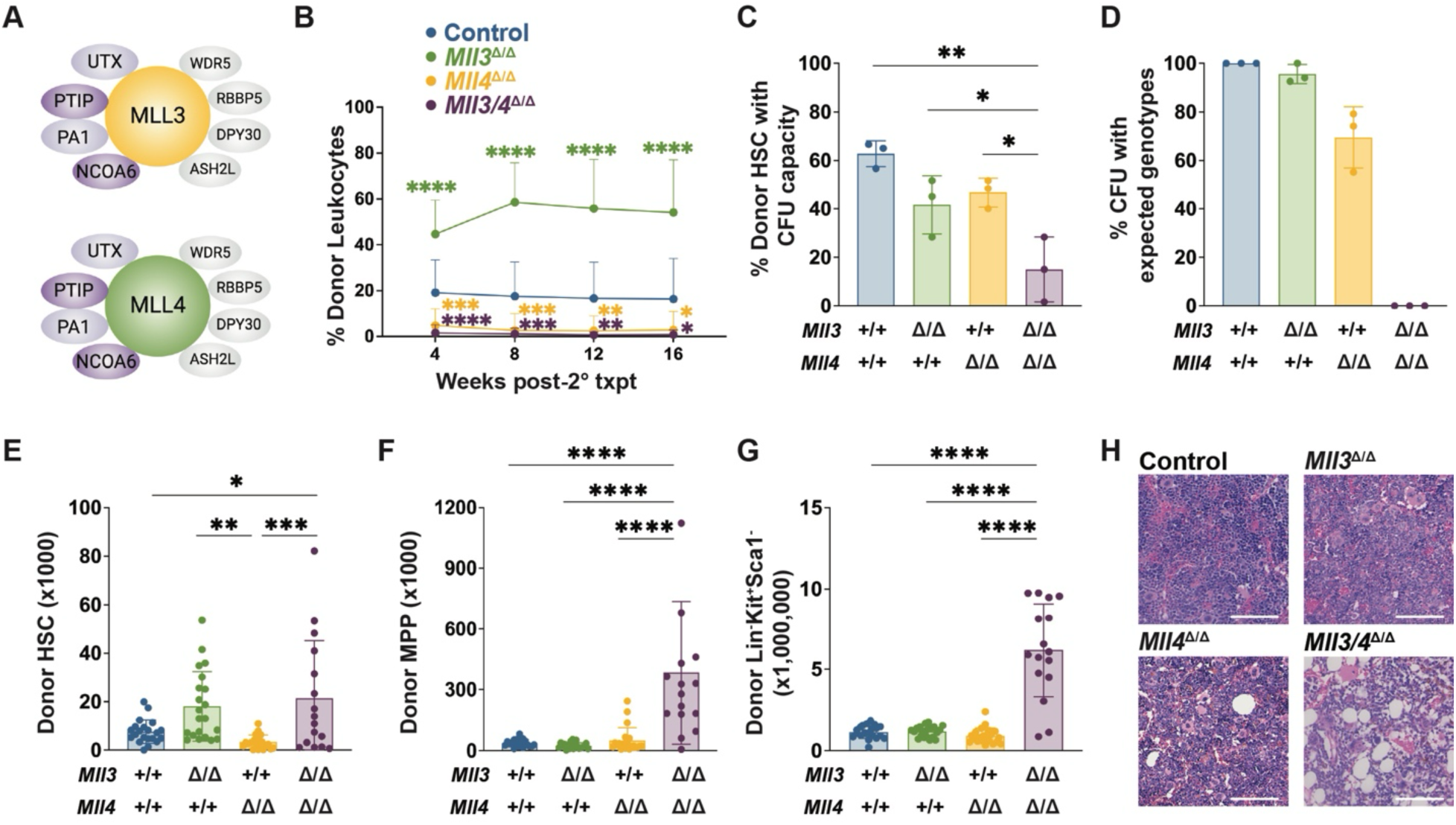
MLL3 and MLL4 act redundantly to sustain myeloid colony formation and hematopoiesis. (A) MLL3- and MLL4-COMPASS complexes. Purple indicates MLL3/4-specific co-factors. (B) Donor (CD45.2^+^) blood chimerism in secondary recipients of indicated donor bone marrow genotypes. (n=15-20). (C) Percentage of donor HSCs with myeloid colony-forming unit (CFU) potential. (n=3). (D) Percentage of myeloid colonies with genotypes that would be expected following complete deletion of floxed alleles. (n=3). (E-G) Donor HSC, MPP, and Lineage^-^Kit^+^Sca1^-^ numbers in recipients at 2 weeks after pIpC treatment (two hindlimbs). (n=16-21). (H) Representative bone marrow histology 30-weeks after pIpC treatment. (n=8-11). Scale bars represent 100µm. For all panels, error bars reflect standard deviation, *p<0.05, **p<0.01, ***p<0.001, ****p<0.0001 by one-way ANOVA and Holm-Sidak posthoc test.

In this study, we conditionally deleted *Mll3*, *Mll4* or both genes together to determine how each gene controls divergent HSC fate decisions. We found that interactions between *Mll3* and *Mll4* are far more complex than a simple antagonism model would suggest, as the genes had unanticipated redundant roles in maintaining the multipotent state. In the absence of both *Mll3* and *Mll4*, hematopoietic progenitors rapidly adopted a non-self-renewing, B-cell-like default identity, coupled with near-total loss of HSC and MPP superenhancer activity. MLL3 and MLL4 proteins did not require catalytically active SET domains to sustain the multipotent state. The data show that myeloid, erythroid and lymphoid potentials are held in tension against one another within HSCs and other progenitors by distinct MLL3/4-dependent and MLL3/4-independent enhancer networks. Our data further illustrate how key functions of MLL3 and MLL4 can be decoupled from the enzymatic functions of each protein.

## Methods

All mouse lines have been previously described.^12,13,25^ For all in vivo assays, cells were isolated, stained, and analyzed as previously described^12,26^ and as discussed in the supplemental methods. CITE-seq, 5’ scRNA-seq, scATAC-seq, ChIPmentation and CUT&RUN libraries were prepared as previously described^26–28^ and analyzed using established computational pipelines.^29–34^ All other methods are described in detail in the Supplemental Methods.

## Results

### MLL3 and MLL4 act redundantly to enable myeloid potential and sustain hematopoiesis

The previously described, opposing phenotypes of *Mll3*- and *Mll4*-deficient HSCs raise the question of whether the encoded proteins interact within a common pathway, such that one gene is epistatic to the other, or whether they have additional redundant functions. To test these possibilities, we competitively transplanted 300,000 *Mx1-Cre*^-^ (Control), *Mll3^f/f^; Mx1-Cre* (*Mll3*^Δ/Δ^), *Mll4^f/f^; Mx1-Cre* (*Mll4*^Δ/Δ^) or *Mll3^f/f^; Mll4^f/f^; Mx1-Cre* (*Mll3/4*^Δ/Δ^) bone marrow cells (CD45.2^+^), along with 300,000 wildtype CD45.1^+^ competitor cells, into lethally irradiated CD45.1^+^ recipient mice (Figure S1A). We confirmed engraftment at 4 weeks post-transplant (Figure S1B), then administered poly-inosine:poly-cytosine (pIpC) to delete the floxed alleles. This approach enabled us to delete *Mll3/4* exclusively in hematopoietic cells without confounding effects of systemic deletion. To assess repopulating activity, we transplanted 3 million bone marrow cells per recipient from the conditionally deleted primary recipient mice into lethally irradiated secondary recipients. *Mll3^Δ/Δ^* marrow engrafted secondary recipient mice more efficiently than control marrow, whereas *Mll4^Δ/Δ^* and *Mll3/4^Δ/Δ^* marrow failed to engraft altogether (Figure 1B; Figure S1C). These phenotypes align with prior studies,^12–14^ and the compound mutant engraftment pattern suggests that *Mll4* may be epistatic to *Mll3* rather than redundant. However, to confirm that *Mll3* and *Mll4* alleles were truly deleted, we genotyped colonies from individual HSCs. Strikingly, none of the *Mll3/4*^Δ/Δ^ HSC-derived colonies had homozygous deletions of both *Mll3* and *Mll4* (Figures 1C-D). Thus, *Mll3* and *Mll4* act redundantly to enable myeloid colony formation from HSCs.

Based on these observations, we tested whether *Mll3* and *Mll4* are necessary to sustain hematopoiesis over time. We noncompetitively transplanted bone marrow from *Mll3/4*^Δ/Δ^ mice (prior to pIpC treatment), administered pIpC 6 weeks after transplantation and analyzed stem and progenitor cell frequencies 2 weeks later (Figures S1D-E). *Mll3/4*^Δ/Δ^ recipients had marked expansion of phenotypic HSCs, MPPs and committed progenitor (Lineage^-^Kit^+^Sca1^-^) populations relative to control recipients (Figures 1E-G; Figures S1F-G). The degree of MPP and committed progenitor expansion was far greater in *Mll3/4*^Δ/Δ^ recipients than in single mutant recipients (Figures 1F-G). We followed a separate cohort for up to 30 weeks post-pIpC, at which point several mice in the *Mll3/4*^Δ/Δ^ recipient cohort became moribund. These mice had pancytopenia, with absolute reductions in all bone marrow lineages and hypocellular morphology (Figure 1H; Figure S2). While *Mll4^Δ/Δ^* recipients also had hypocellular marrow (Figure S2E-I), they did not have aberrant expansion of HSC, MPP or committed progenitor populations, suggesting that cytopenias in these mice reflect progenitor attrition rather than a differentiation block. Altogether, the changes in phenotypic progenitor numbers observed in *Mll3/4*^Δ/Δ^ recipients suggest redundant roles for MLL3 and MLL4 in HSC/MPP fate specification, in addition to their non-redundant, antagonistic functions.

### MLL3 and MLL4 maintain multilineage hematopoiesis by opposing a B-cell-like default state

To better evaluate the molecular changes that ensue following compound *Mll3/4* deletion, we performed Cellular Indexing of Transcripts and Epitopes by Sequencing (CITE-seq) on donor Lineage^-^Kit^+^Sca1^+^ (LSK) cells as well as Kit^+^ cells isolated from primary recipient bone marrow two weeks after pIpC treatment (Figure S1D; Table S1). Iterative Clustering with Guide-gene Selection (ICGS) identified 20 transcriptionally distinct cell clusters across all genotypes (Figures 2A-B; Figure S3A). In some cases, clusters from adjacent clades of the ICGS heatmap were combined as superclusters (called HSC, MPP4, MLL4 ko-1, MLL4 ko-2 and DKO) to simplify description of the data (Figures 2A-B; Figure S3A). We identified populations within each sample group with previously described HSC/MPP immunophenotypes (Figure 2C; Figures S3B-C).^5^ Control and *Mll3*^Δ/Δ^ LSK had similar cluster distributions, with modest expansion of the HSC supercluster in *Mll3*^Δ/Δ^ LSK (Figure 2D). *Mll4*^Δ/Δ^ and *Mll3/4*^Δ/Δ^ LSK had very different distributions than controls despite having HSC and MPP immunophenotypes (Figures 2B-C). *Mll4*^Δ/Δ^ LSK populated two superclusters (MLL4 ko-1 and 2) and ectopically expressed myeloid genes, including *Elane*, *Mpo*, and *Il1rl1* (Figure 2E; Figure S3D). *Mll3/4*^Δ/Δ^ LSK populated a distinct supercluster (DKO) and ectopically expressed B-cell genes, including *Il7r*, *Dntt*, *Pax5*, and *Ebf1* (Figure 2F; Figure S3E; Table S2). Thus, *Mll3* limits the size of the HSC pool, *Mll4* prevents precocious myeloid gene expression, and *Mll3* and *Mll4* act redundantly to prevent conversion of HSC/MPPs to a B-cell-like state.

**Figure 2.**
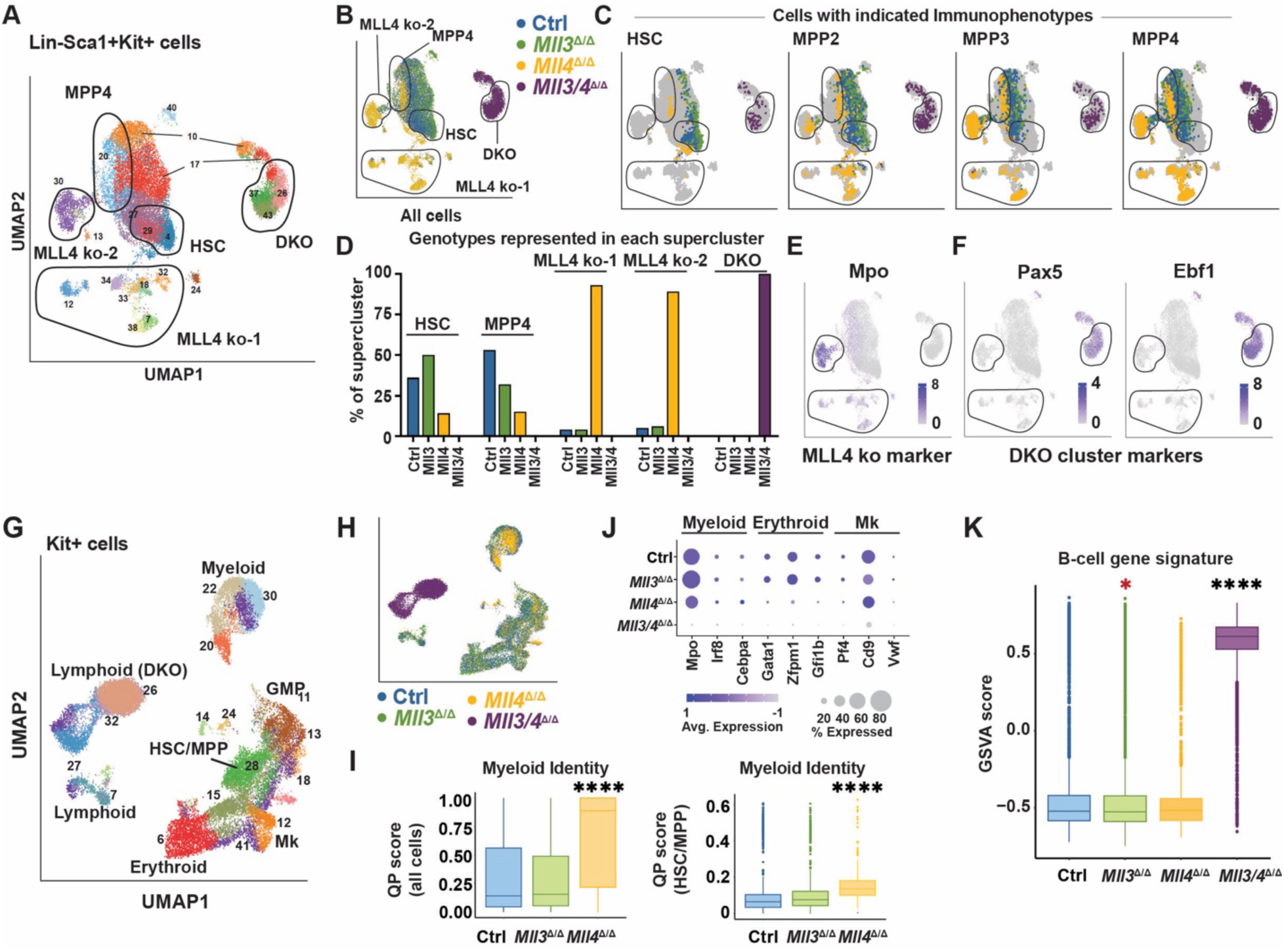
Compound *Mll*3/4 deletion converts hematopoietic progenitors to a B-cell-like state. (A, B) UMAP representing donor LSK single cell transcriptomes, colored by cluster (A) or genotype (B). Five superclusters are outlined in black. (C) Distribution of HSC, MPP2, MPP3, and MPP4 immunophenotypes by genotype. (D) HSC, MPP2, MPP3 and MPP4 immunophenotype frequencies in each supercluster. (E, F) Expression of myeloid and lymphoid marker genes in single cells. (G, H) UMAPs representing donor Kit^+^ single cell transcriptomes, colored by cluster (G) or genotype (H). HSC/MPP, GMP, myeloid, lymphoid, erythroid, and megakaryocytic (Mk) annotations are indicated. (I) Myeloid identity quadratic programming (QP) scores for HSC/MPP, GMP and myeloid superclusters combined (left) and the HSC/MPP cluster alone (right). ****p<0.0001 for *Mll4*^Δ/Δ^ relative to control or *Mll3*^Δ/Δ^ by pairwise Wilcoxon test. (J) Myeloid, erythroid, and Mk marker gene expression by genotype. (K) B-cell GSVA enrichment scores by genotype. ****p<0.0001 in *Mll3/4*^Δ/Δ^ (higher) relative to all other genotypes, *p=0.02 in *Mll3*^Δ/Δ^ (lower) relative to control.

We next performed CITE-seq on donor Kit^+^ cells to capture effects of *Mll3* and *Mll4* deletions on more differentiated progenitors. We annotated clusters that contained HSC/MPP, granulocyte-monocyte progenitor (GMP), myeloid, lymphoid, erythroid, and megakaryocytic cells based on gene and surface marker expression (Figures 2G-H; Figures S4A-B). *Mll3* deletion had only modest effects on the cluster distributions (Figure S4C). *Mll4* deletion increased myeloid cluster representation and reduced HSC/MPP and GMP cluster representation (Figure S4C). To more precisely test for enhanced myeloid bias in *Mll4*^Δ/Δ^ progenitors, we used quadratic programming to calculate myeloid identity scores for each cell within the HSC/MPP, GMP and myeloid clusters. *Mll4* deletion significantly increased myeloid identity scores across all myeloid progenitor populations, including HSC/MPPs (Figure 2I; Figure S4D), consistent with precocious myeloid differentiation. In contrast, *Mll3/4*^Δ/Δ^ cells clustered separately from all other cells (DKO cluster) and expressed B-cell genes, including *Pax5* and *Ebf1* (Figure S4E). The cells also expressed CD127, and most had a common lymphoid progenitor (CLP) surface marker phenotype (Figures S4A-B). There was no evidence of myeloid, erythroid or megakaryocyte gene expression in *Mll3/4*^Δ/Δ^ cells (Figure 2J). Instead, Gene Set Variant Analysis (GSVA) confirmed strong enrichment of a B-cell signature (Table S3) within most *Mll3/4*^Δ/Δ^ cells (Figure 2K). Similar changes progenitor numbers, function and gene expression were observed when Ubc-CreER was used to delete *Mll3* and *Mll4*, indicating that they were not caused by pIpC (Figures S5A-I). Deleting *Mll3* and *Mll4* in committed myeloid progenitors, with *LysM*-Cre, did not impair terminal myelopoiesis or cause B-cell reprogramming (Figures S5J-M). Thus, once myeloid commitment is fully established, *Mll3* and *Mll4* become dispensable for later stages of myeloid differentiation. These data show that MLL3 and MLL4 act redundantly within HSC/MPPs to maintain myeloid, erythroid and megakaryocytic potential, and they oppose a B-cell-like default state.

### Mll3/4-deficient progenitors do not resemble normal B-cell progenitors

We next sought to characterize the B-cell-like, *Mll3/4*-deficient progenitor population in greater detail, with the goal of understanding whether the population aligns with a stage of normal B lymphopoiesis or whether it reflects an emergent cell state that lacks a clear analog in normal hematopoiesis. We tested whether *Mll3/4*^Δ/Δ^ progenitors have B-cell potential by plating control, *Mll3*^Δ/Δ^, *Mll4*^Δ/Δ^ and *Mll3/4*^Δ/Δ^ LSK cells at limiting dilutions and culturing in conditions that support B-cell production. *Mll3* deletion enhanced B-cell colony formation, *Mll4* deletion impaired colony formation, and despite B-cell priming, *Mll3/4*^Δ/Δ^ LSK lacked B-cell colony forming potential altogether (Figures 3A-B). Furthermore, *Mll3/4*^Δ/Δ^ mice had reduced donor pro-B, pre-B, and B220^hi^ recirculating B-cell numbers in the bone marrow (Figure 3C; Figure S6A), indicating B-cell maturation defects.

**Figure 3.**
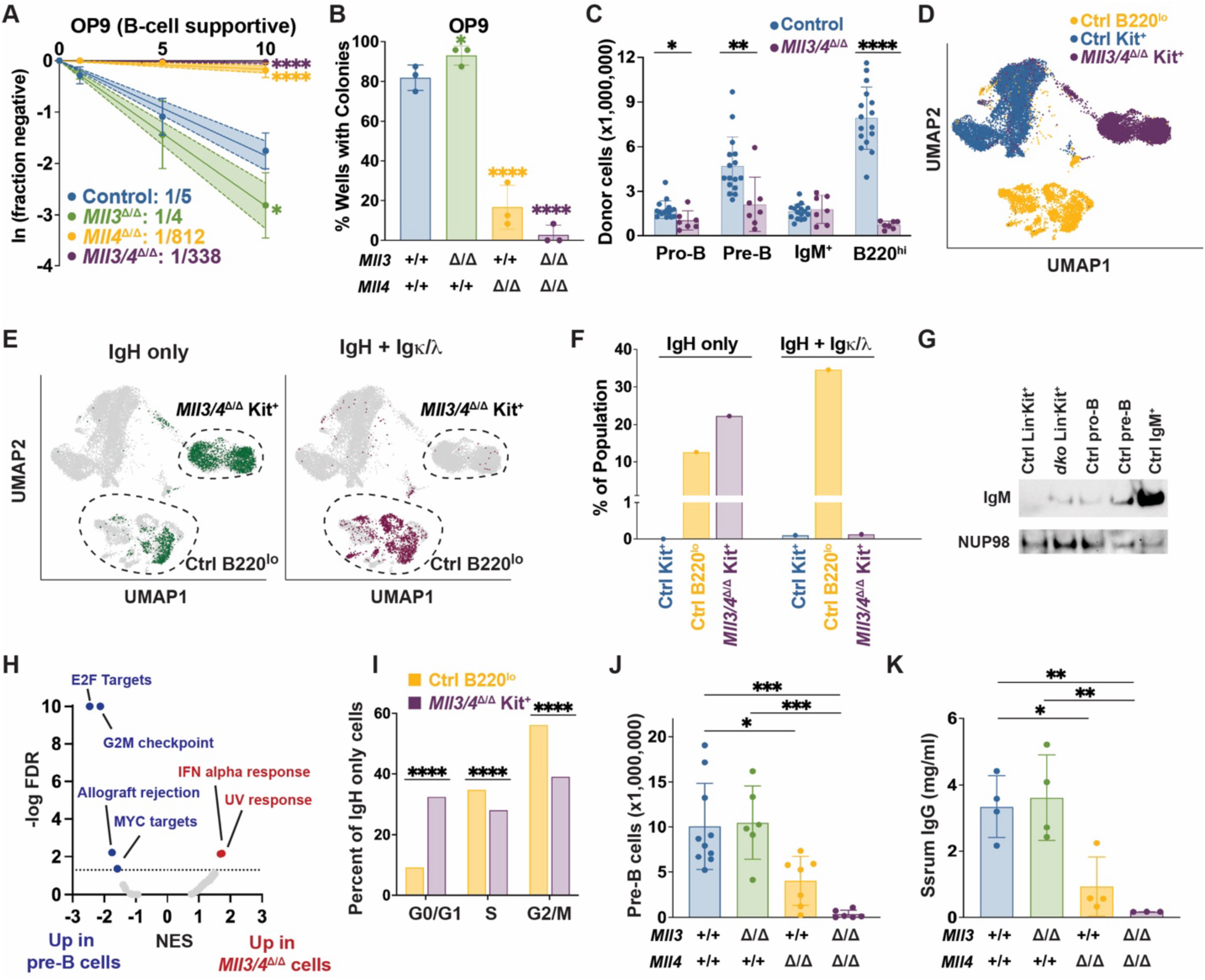
*Mll3/4* deficient HSC/MPPs lack mature B-cell potential and undergo immunoglobin heavy but not light chain rearrangement. (A) Limiting dilution analysis of B-cell colony forming potential of donor LSK cells with indicated genotypes. (n=3). (B) Percent of wells with B-cell colonies when 10 donor cells/well were plated. *p<0.05, ****p<0.0001 relative to control, by ELDA. (C) Donor pro-B, pre-B, IgM^+^, and B220^hi^ recirculating B-cells in the bone marrow of control or *Mll3/4*^Δ/Δ^ recipient mice. (n=7-16). (D) Distribution of single cell transcriptomes for control and *Mll3/4*^Δ/Δ^ genotypes. Donor control B220^lo^ and Kit^+^ cells serve as positive and negative controls for immunoglobin rearrangement, respectively. (E) Distributions of cells with heavy chain rearrangements only (left, IgH) or heavy and light chain rearrangements (right, IgH + Igξ/1). Distinct clusters of control B220^lo^ cells and *Mll3/4*^Δ/Δ^ Kit^+^ cells are circled. (F) Percentage of cells with IgH only or both IgH + Igξ/1 rearrangement by cell type. (G) IgM expression in the indicated cell populations and genotypes. NUP98 serves as a loading control. (H) Volcano plot showing differentially enriched Hallmark gene sets between IgH-rearranged control B220^+^ and *Mll3/4*^Δ/Δ^ Kit^+^ cells. (I) Calculated cell cycle phases for IgH-rearranged cells. ****p<0.0001 by Chi-square test. (J) Bone marrow pre-B cells in mice of indicated genotypes following deletion with *Mb1*-Cre. (n=6-10). (K) Serum IgG levels in the *MBI-Cre* conditional knockout mice. (n=3-4). Error bars reflect standard deviation, *p<0.05, **p<0.01, ***p<0.001, ****p<0.0001 by one-way ANOVA followed by Holm-Sidak posthoc test.

We next used single cell RNA-sequencing to test whether *Mll3/4*^Δ/Δ^ cells are capable of rearranging immunoglobin (*Ig*) loci. For these assays, we included control and *Mll3/4*^Δ/Δ^ Kit^+^ cells, as well as Cre-negative B220^lo^ bone marrow cells. *Mll3/4*^Δ/Δ^ cells clustered separately from both control Kit^+^ and B220^lo^ lymphoid progenitor cells (Figure 3D). They underwent *Ig* heavy chain (IgH) rearrangement without further light chain rearrangement, in contrast to control B220^lo^ cells (Figures 3E-F). Western blotting for IgM showed low levels of protein expression in the *Mll3/4*^Δ/Δ^ cells (Figure 3G), likely reflecting IgH but not mature IgM expression. Gene set enrichment analysis (GSEA) with pseudobulk populations revealed several differences between control and *Mll3/4*^Δ/Δ^ IgH-rearranged cells, including evidence of reduced proliferation within the *Mll3/4*^Δ/Δ^ population (Figures 3H-I). Altogether, the data show that the aberrant, B-cell-like default state of *Mll3/4*^Δ/Δ^ progenitors lacks a clear analog within the normal B-cell differentiation trajectory.

We next tested whether deleting *Mll3 and Mll4* in pro-B-cells, using *Mb1-Cre*, would impede B-cell maturation. *Mll4* deletion significantly reduced pro-B, pre-B, and B220^hi^ recirculating B-cells in the bone marrow, as well as follicular, marginal zone, and transitional B-cell populations in the spleen (Figure 3J; Figures S6B-I). *Mll4* deficiency also reduced serum IgG levels, consistent with loss of functional mature B-cells (Figure 3K). Compound *Mll3/4* deletions appeared to exacerbate these phenotypes, though the changes were not statistically significant. The data illustrate stage-specific roles for MLL3 and MLL4 during B-cell development. B-cell priming in HSC/MPPs occurs independently of MLL3/4, whereas B-cell maturation requires MLL4, with modest redundancy afforded by MLL3.

### MLL4 maintains the HSC/MPP enhancer landscape by preventing precocious activation of myeloid enhancers

To understand how MLL3 and MLL4 regulate enhancer activity to maintain HSC/MPP multipotency, restrict myeloid differentiation and prevent ectopic B-cell priming, we performed single cell ATAC-seq (scATAC-seq) on control, *Mll3*^Δ/Δ^, *Mll4*^Δ/Δ^ and *Mll3/4*^Δ/Δ^ LSK cells (Figure S1D). We clustered cells in ArchR^30^ and used integrated CITE-seq data to annotate the clusters based on inferred transcript expression (Figures 4A-C). We identified HSC, MPP4, MLL4 ko-1, MLL4 ko-2 and DKO superclusters, analogous to the superclusters defined in Figures 2A and S3A, based on integrated gene expression (Figure S7A-B). As anticipated, control and *Mll3*^Δ/Δ^ cells had very similar cluster distributions (Figure 4A; Figure S7B). In contrast, *Mll4*^Δ/Δ^ cells occupied MLL4 ko-1 and MLL4 ko-2 superclusters, and *Mll3/4*^Δ/Δ^ cells occupied the DKO cluster almost exclusively (Figures S7B-C). These scATAC-seq profiles recapitulate cluster distributions observed by CITE-seq.

**Figure 4.**
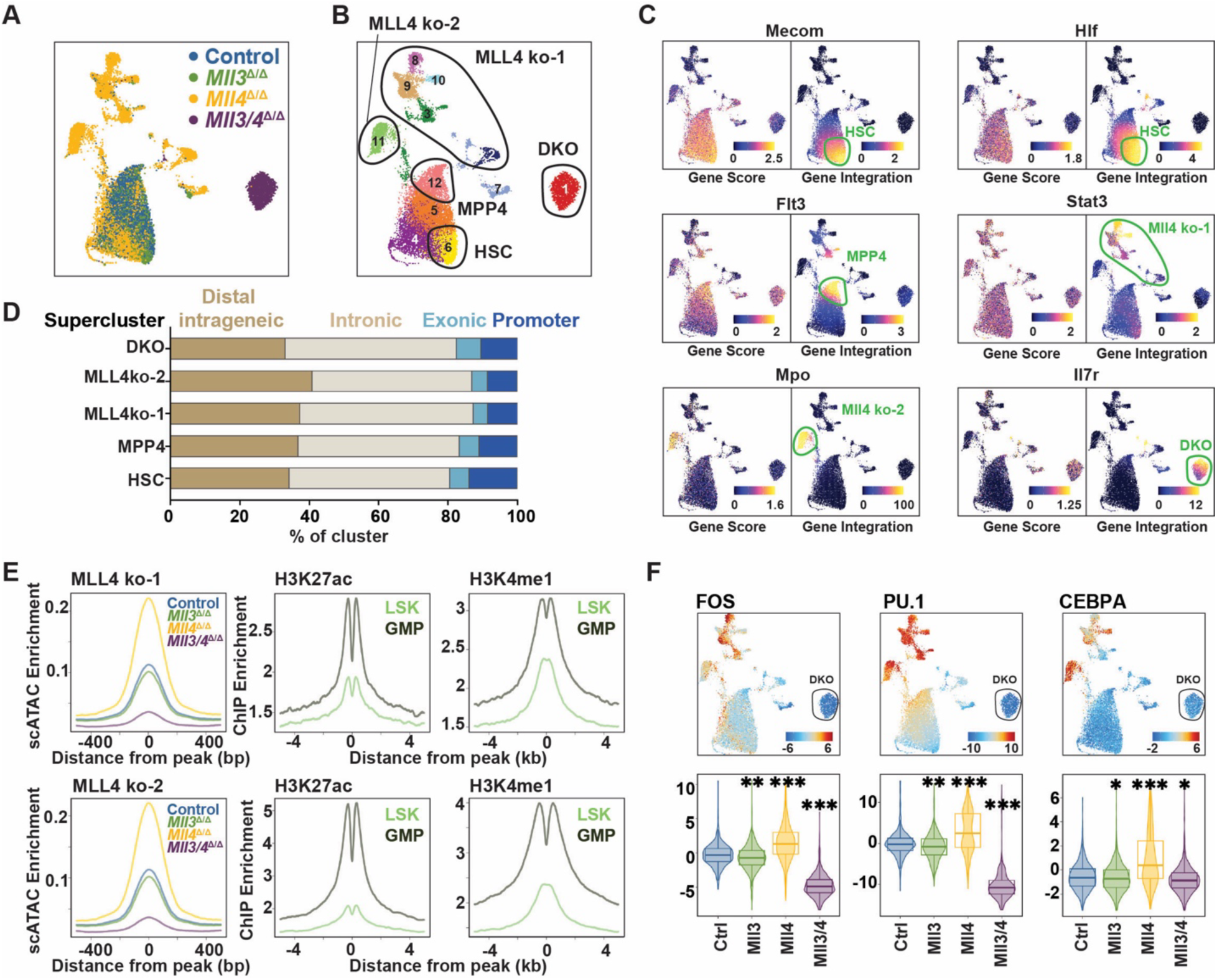
*Mll4* deletion causes premature activation of GMP enhancers in HSC/MPPs. (A, B) scATAC-seq profiles of donor LSK, by genotype or ArchR clusters, with superclusters annotation based on integrated CITE-seq data. (C) Accessibility and integrated expression of representative HSC, MPP, myeloid and B-lymphoid marker genes. (D) Distribution of scATAC-seq peaks relative to transcriptional start sites by supercluster. (E) Left: Supercluster-specific aggregate scATAC-seq signals by genotype. Middle and right: H3K27ac and H3K4me1 signals at MLL4-ko-1 and MLL4-ko-2-specific elements, for LSK or GMP. (F) ChromVAR deviation scores for FOS, PU.1, and CEBPA motifs. For all violin plots, *p<0.05, **p<10^-10^, ***p<10^-100^, by Wilcoxon signed rank test relative to control.

Cluster-specific, differentially accessible regions mapped primarily to distal enhancer elements. Approximately 80-90% of the elements were located in intragenic or intronic regions (Figure 4D; Table S4). HSC and MPP4 cluster-specific elements overlapped with previously curated H3K4me1 and H3K27ac peaks from sorted HSCs and MPPs (Figure S7D).^12^ These elements therefore reflect active HSC/MPP enhancers. *Mll3* deletions did not alter H3K4me1 or H3K27ac levels at HSC- or MPP4-specific enhancers (Figure S7E), consistent with a prior study showing that MLL3 regulates H3K4me1 and H3K27ac at only a small number of target enhancers that primarily associate with inflammatory responses.^12^ *Mll4*^Δ/Δ^ LSK enhancers had H3K4me1 and H3K27ac patterns that more closely resembled GMPs than HSC/MPP (Figure 4E), and transcription factor motif enrichment (e.g., FOS, PU.1, CEBPA) that indicated precocious myeloid differentiation (Figure 4F; Figures S7F-G; Table S5). Thus, MLL4 maintains the HSC/MPP enhancer landscape and prevents premature activation of GMP-associated enhancers.

### MLL3 and MLL4 act redundantly to maintain HSC/MPP enhancers while suppressing activity of B-cell regulatory elements

As in the CITE-seq studies, *Mll3/4*^Δ/Δ^ LSK clustered apart from all normal HSCs and MPPs based on their scATAC-seq profiles (DKO supercluster; Figures 4A-B). We confirmed that cells in the DKO supercluster have increased accessibility of B-cell-associated genes by GSVA (Figure S8A) and normal expression of other COMPASS proteins by Western blot (Figure S8B). DKO-specific elements were enriched for EBF1 and PAX5 motifs (Figures 5A-B; Figure S8C), consistent with B-cell differentiation.^35^ Analysis of previously-described pro-B-cell ChIP-seq data confirmed binding of EBF1^36^ and PAX5^37^ to DKO-specific enhancers (Figure 5C). Motifs for critical effectors of HSC and myeloid identity (e.g., NFIX, RUNX1 and GATA2) were negatively enriched in DKO-specific enhancers (Figure 5D; Figure S8C). We performed ChIPmentation on wildtype and *Mll3/4*^Δ/Δ^ LSK cells to evaluate H3K27ac (a mark of active enhancers and promoters) at HSC/MPP cis-regulatory elements. *Mll3*/*4* deletion did not alter total H3K27ac levels in LSK, based on Western blot (Figures S8D-E), but it dramatically reduced H3K27ac at a previously curated set of enhancers that are normally active in HSC/MPPs (Figure 5E).^12^ This reduction was offset by gain of H3K27ac at DKO-specific enhancers (Figure 5F). In addition, the promoters and gene bodies of several B-cell regulators, including *Ebf1* and *Pax5*, were hyperacetylated in *Mll3/4*^Δ/Δ^ LSK (Figures 5G-H). Thus, *Mll3/4* deletion causes dramatic realignment of HSC/MPP enhancer activities, such that enhancers that maintain stemness and myeloid identity are inactivated while enhancers that promote B-cell differentiation are ectopically activated.

**Figure 5.**
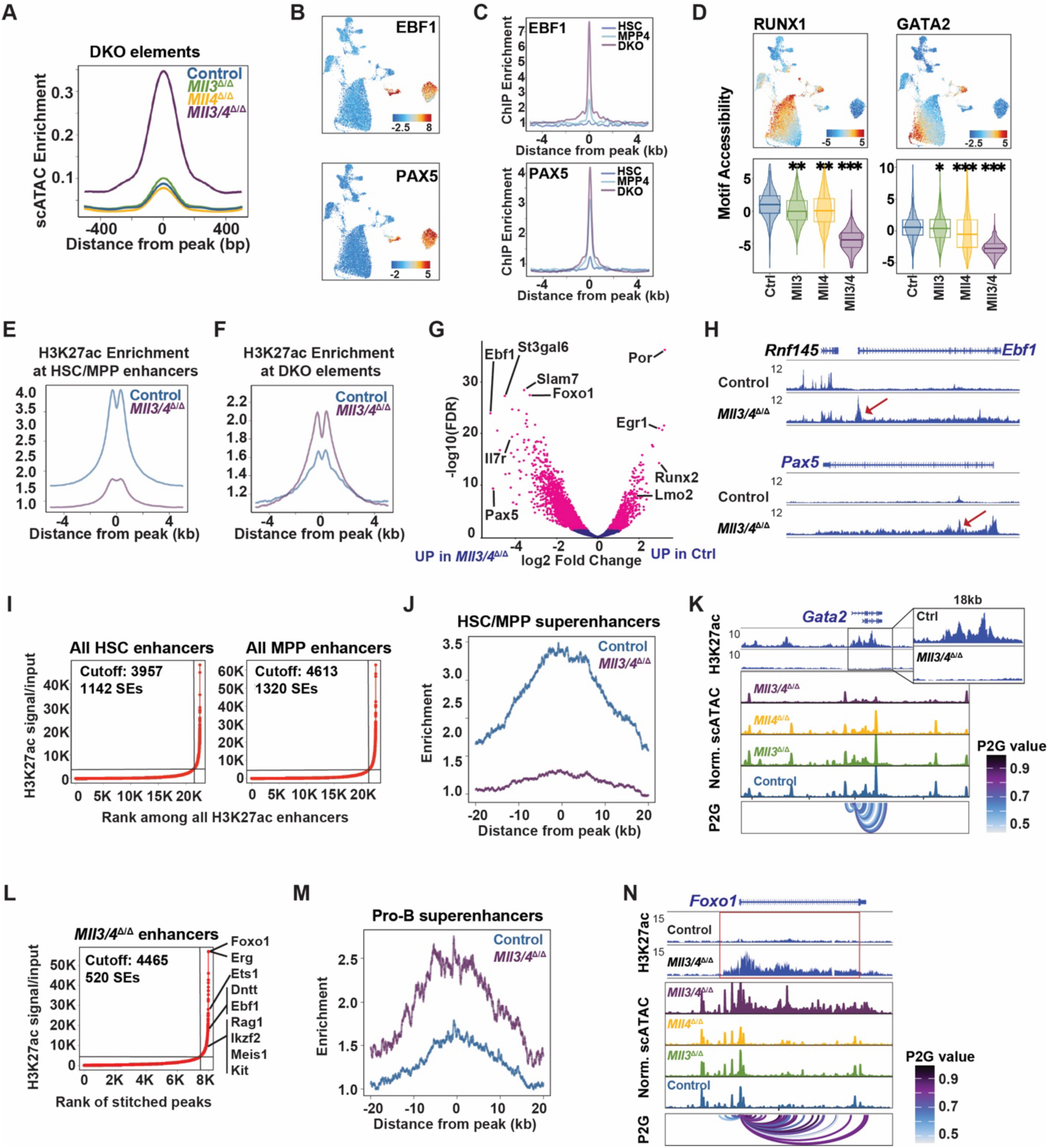
Loss of MLL3 and MLL4 inactivates HSC/MPP enhancer networks and ectopically activates B-cell enhancers. (A) DKO cluster-specific aggregate scATAC-seq signals by genotype. (B) ChromVAR deviation for EBF1 and PAX5 motif accessibility. (C) EBF1 and PAX5 binding in pro-B cells at cluster-specific enhancers. (D) ChromVAR deviation for RUNX1 and GATA2 showing reduced accessibility in *Mll3/4*^Δ/Δ^ LSK. *p<0.05, **p<10^-10^, ***p<10^-100^, by Wilcoxon tests. (E, F) H3K27ac signal at HSC/MPP and DKO-specific enhancers in control and *Mll3/4*^Δ/Δ^ LSK. (G) Volcano plot showing differential H3K27ac at DKO-specific enhancers in control and *Mll3/4*^Δ/Δ^ LSK cells. (H) H3K27ac tracks at *Ebf1* and *Pax5* loci. Red arrows indicate regions of significantly increased H3K27ac in *Mll3/4*^Δ/Δ^ LSK. (I) Rank Ordering of Super Enhancers (ROSE) analysis of HSC and MPP enhancers based on H3K27ac signals. (J) Aggregate H3K27ac levels at HSC/MPP superenhancers in control and *Mll3/4*^Δ/Δ^ LSK cells. (K) Tracks showing reduced accessibility and H3K27ac at *Gata2* in *Mll3/4*^Δ/Δ^ LSK. Peaks to gene (P2G) indicates association between peaks and the promoter. (L) ROSE analysis to identify MLL3/4-independent superenhancers in *Mll3/4*^Δ/Δ^ cells. (M) Aggregate H3K27ac levels at previously described pro-B-cell superenhancers in control and *Mll3/4*^Δ/Δ^ LSK. (N) Tracks showing increased accessibility and H3K27ac at *Foxo1* in *Mll3/4*^Δ/Δ^ LSK. Red boxes indicate MLL3/4-independent superenhancers.

The transition from HSC/MPP to B-cell regulatory programs was even more pronounced when we evaluated superenhancers. Superenhancers encompass large genomic regions and are notable for very high H3K27ac levels. We identified HSC and MPP superenhancers by performing Rank Order of Super Enhancers (ROSE)^33^ and then evaluated H3K27ac levels in *Mll3/4*^Δ/Δ^ LSK (Figure 5I; Table S6).^12^ *Mll3/4* deletion dramatically reduced H3K27ac in these regions, including at superenhancers associated with critical HSC genes such as *Myct1*, *Gata2*, *Mecom*, *Cd34*, and *Hlf* (Figures 5J-K; Figures S8F-G). These reductions in superenhancer activity were accompanied by activation of pro-B-cell associated superenhancers in *Mll3/4*^Δ/Δ^ LSK,^33^ including at genes such as *Ebf1*, *Foxo1*, *Rag1*, *Dntt* and *Il7r* (Figures 5L-N; Figure S8H-I). Thus, MLL3 and MLL4 are necessary to maintain normal HSC/MPP superenhancer activity and suppress B-cell superenhancer activity.

### MLL3/4 directly bind enhancers near HSC/MPP transcription factor genes but not B-cell-specific genes

Single cell transcript expression and chromatin profiling data predict that MLL3 and MLL4 directly and redundantly regulate expression of transcription factors that maintain HSC self-renewal and multipotency. To identify and characterize MLL3/4-bound regulatory elements, we performed CUT&RUN using antibodies directed at MLL3 or MLL4. In parallel, we engineered mouse 32D cells to express an N-terminal 3xFLAG-tagged MLL3 protein from both *Mll3* alleles to facilitate ChIPmentation, as well as an *Mll3*-null negative control line (Figure 6A). Co-immunoprecipitation assays confirmed expression of 3xFLAG-MLL3 and association with COMPASS co-factors (Figure S9A). CUT&RUN identified far more MLL3 bound regions than ChIPmentation with a FLAG-specific antibody, suggesting greater sensitivity. Essentially all FLAG-bound sites (96%) overlapped with both MLL3 and MLL4 CUT&RUN peaks (Figure 6B). We focused on these elements, even though they represent only a small fraction of the overlapping MLL3 and MLL4 CUT&RUN peaks, because they were supported by orthogonal assays. Most MLL3/4-bound elements mapped to distal intragenic or intronic regions, and ∼75% of all overlapping peaks aligned with previously described HSC/MPP enhancers,^12^ GMP enhancers,^26^ or both (Figures 6C-D). MLL3/4-bound regions overlapped with UTX binding sites,^38^ as expected for a COMPASS protein (Figure 6E), and were enriched for ETS, RUNX and STAT5 binding motifs (Figure S9B). Deleting *Mll3* in HSCs, MPPs, and GMPs had only modest effects on H3K27ac at MLL3/4-bound regions (Figure 6F; Figure S9C). In contrast, deleting *Mll3/4* together largely attenuated H3K27ac at those regions, indicating redundancy between MLL3 and MLL4 (Figure 6F). Many MLL3/4-bound enhancers mapped near genes that sustain HSC self-renewal, including *Runx1*, *Cited2*, *Cux1, Zeb2* and *Erg* (Figures 6G-H; Figure S9D; Table S7). They did not map near genes that prime B-cell identity, such as *Ebf1* or *Pax5*, nor did they map near other B-cell identity genes (e.g, *Cd19*, *Cd79a*, *Dntt*, etc.). We observed minimal overlap with DKO cluster-specific regulatory elements (8 out of 1013 total elements). These data imply that MLL3 and MLL4 directly and redundantly regulate expression of transcription factors that sustain HSC/MPP multipotency. In doing so, they indirectly repress genes that prime B-cell identity.

**Figure 6.**
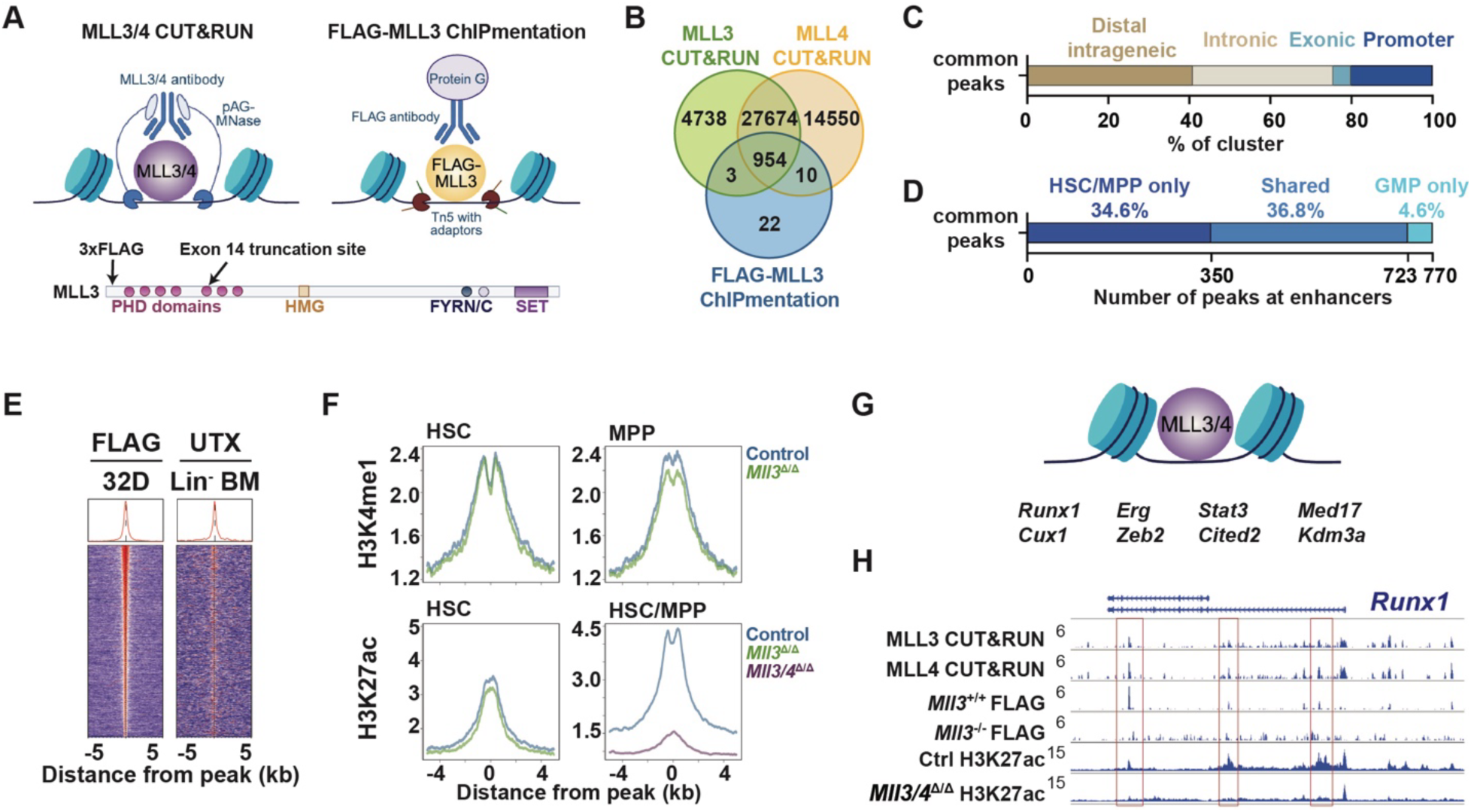
MLL3 and MLL4 directly and redundantly regulate a subset of HSC/MPP transcription factors. (A) Top: Overview of CUT&RUN and ChIPmentation strategies as orthogonal methods to study MLL3/4 chromatin localization. Bottom: Schematic of 3xFLAG-MLL3 insertion, and location of the inactivating exon 14 mutations, in 3xFLAG-MLL3 and 3xFLAG-MLL3-null 32D cells, respectively. (B) Venn diagram showing overlapping peaks among MLL3 CUT&RUN, MLL4 CUT&RUN, and FLAG-MLL3 ChIPmentation datasets. (C) Distribution of MLL3/4-bound peaks (the intersection in Figure 6B) relative to gene bodies. (D) Association of MLL3/4-bound peaks with enhancers that are either primed (H3K4me1) or active (H3K4me1 and H3K27ac) in HSC/MPPs, GMPs or both. (E) Tornado plots showing UTX binding in Lineage^-^ bone marrow at MLL3/4-bound enhancers. FLAG-MLL3 signal is shown to the left. (F) Top: H3K4me1 at MLL3/4-bound enhancers in control and *Mll3*^Δ/Δ^ HSCs and MPPs. Bottom: H3K27ac at MLL3/4-bound enhancers in control and *Mll3*^Δ/Δ^ HSCs, and control or *Mll3/4*^Δ/Δ^ LSK. (G) Schematic of MLL3 and MLL4 target genes encoding key HSC/MPP transcription factors or epigenetic regulators. (H) Representative tracks showing MLL3/4 CUT&RUN, FLAG-MLL3 ChIPmentation, and H3K27ac at the *Runx1* locus. Red boxes indicate MLL3/4-bound enhancers.

### MLL3 and MLL4 sustain multilineage hematopoiesis independently of histone methyltransferase activity

We next tested whether MLL3 and MLL4 require histone methyltransferase activity, mediated by the SET domains in each protein, to sustain HSC/MPP multipotency. We crossed previously described *Mll3^Y4792A^* and *Mll4^Y5477A^* mutant alleles^25^ with *Mll3^f/f^; Mll4^f/f^; Mx1-Cre* mice to generate Cre-negative (Control), *Mll3^Y4792A/f^; Mx1-Cre* (*Mll3*^SI^)*, Mll4^Y5477A/f^; Mx1-Cre* (*Mll4*^SI^) and *Mll3^Y4792A/f^; Mll4^Y5477A/f^; Mx1-Cre* (*Mll3/4*^SI^) mice. We transplanted bone marrow from control, *Mll3*^SI^, *Mll4*^SI^ and *Mll3/4*^SI^ mice and treated with pIpC to delete the floxed alleles (Figures 7A-B). After pIpC treatment, hematopoietic cells retained single copies of the SET-inactive alleles. SET-inactive LSK cells had reduced H3K4me1 levels that resembled complete MLL3/4 loss of function (Figure 7C). Phenotypic MPP and myeloid progenitor populations expanded, modestly, in *Mll3/4*^SI^ recipients, but the magnitude was lower than was observed in *Mll3/4*^Δ/Δ^ recipients (Figures 7D-F). *Mll3/4*^SI^ HSCs had reduced but not absent myeloid colony forming potential, in contrast to *Mll3/4*^Δ/Δ^ LSK (Figure 7G). Thus, myeloid colony formation does not absolutely require MLL3/4 SET activity.

**Figure 7.**
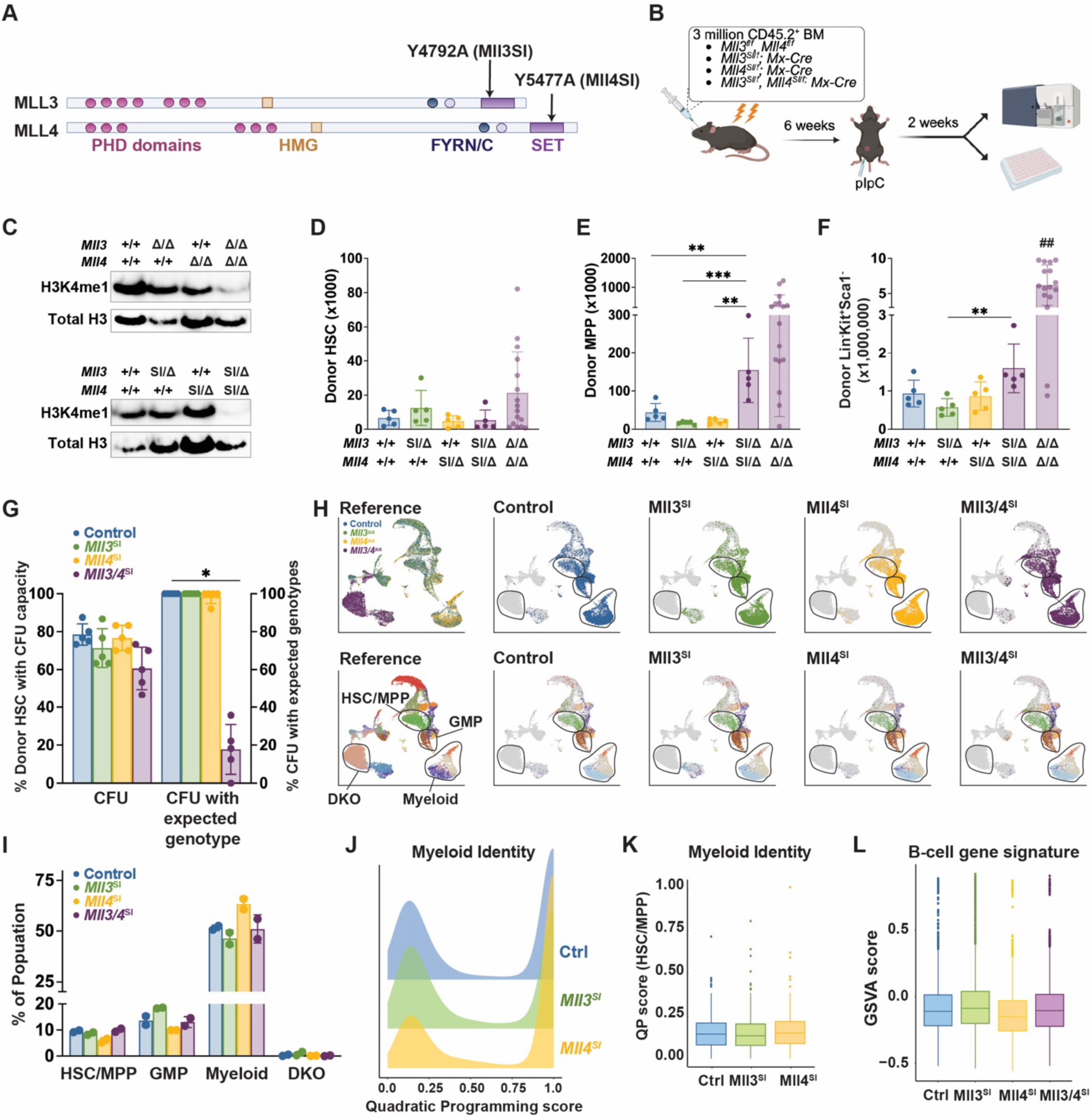
MLL3 and MLL4 do not require histone methyltransferase activity to sustain HSC/MPP multipotency, prevent premature myeloid differentiation or prevent conversion to a B-cell like default state. (A) Schematics indicating MLL3 and MLL4 SET inactivating (SI) mutations. (B) Overview of noncompetitive transplantation strategy to assess dependence on MLL3/4 histone methyltransferase activity. (C) Western blots showing H3K4me1 in LSK cells of indicated genotypes. The SET-inactivating alleles greatly diminished total H3K4me1. (D-F) Donor HSC, MPP, and Lineage^-^Kit^+^Sca1^-^ numbers in recipient bone marrow 2 weeks after pIpC treatment. (n=5). *Mll3/4*^Δ/Δ^ cell numbers from Figure 1 were included for comparison. Error bars reflect standard deviation, *p<0.05, **p<0.01, ***p<0.001 by one-way ANOVA and Holm-Sidak posthoc test, ^##^p<0.01 for comparison of *Mll3/4*^Δ/Δ^ and *Mll3/4*^SI^ by two-tailed Student’s t-test. (G) Percentage of donor HSCs with myeloid colony-forming unit (CFU) potential (left) and percentage of myeloid colonies with genotypes that are expected following complete deletion of floxed alleles (right). (n=3). (H) Distribution of single cell transcriptomes of *Mll3/4^SI^* Kit^+^ cells projected against *Mll3/4*^Δ/Δ^ Kit^+^ cells from Figure 2G by Symphony. Cells are colored by genotype (top) or reference cluster identity (bottom) as in Figure 2G. (I) Percentage of cells for each genotype that map to HSC/MPP, GMP, myeloid and DKO superclusters by Symphony. (n=2). (J, K) Myeloid identity quadratic programming (QP) scores for HSC/MPP, GMP and myeloid clusters combined (J) or the HSC/MPP cluster alone (K). (L) B-cell signature GSVA enrichment scores by genotypes. For J-L, myeloid identity and B-cell enrichment scores did not significantly differ by genotype.

We performed CITE-seq on control, *Mll3*^SI^, *Mll4*^SI^ and *Mll3/4*^SI^ Kit^+^ cells, as in Figure 2G, to test whether MLL4 SET inactivation causes precocious myeloid differentiation and whether MLL3/4 SET inactivation drives cells toward a B-cell-like state, as was observed with complete loss-of-function. For these comparisons, we used Symphony^32^ to assign SET-inactive cells to the same clusters as were defined using the complete loss-of-function dataset in Figure 2G, with the data from Figure 2G serving as the reference. We then performed quadratic programming and GSVA to assess myeloid differentiation and B-cell priming, respectively. In contrast to *Mll4*^Δ/Δ^ cells, *Mll4*^SI^ cells were not enriched in the myeloid supercluster and did not show evidence of precocious myeloid differentiation based on quadratic programming (Figures 7H-K). Likewise, in contrast to *Mll3/4*^Δ/Δ^ cells, *Mll3/4*^SI^ cells did not associate with the DKO supercluster or show evidence of ectopic B-cell gene expression based on GSVA analysis (Figures 7H-I, L). Altogether, these data show that MLL3 and MLL4 maintain HSC/MPP multipotency and prevent precocious myeloid differentiation independently of their histone methyltransferase activities.

## Discussion

This study illuminates critical, redundant roles for MLL3 and MLL4 in maintaining HSC/MPP multipotency and enabling multilineage hematopoiesis. The consequences of MLL3/4 inactivation are far more pervasive than is typically observed for individual transcription factors or epigenetic regulators. For example, loss of transcription factors such as RUNX1, GATA2 or MECOM can impair HSC self-renewal, disrupt quiescence, alter gene expression, and cause attrition of the HSC pool, but populations of myeloid, erythroid and megakaryocytic progenitors can still be identified in the absence of these genes.^39–41^ Likewise, loss of epigenetic regulators, such as DNMT3A, TET2 or ASXL1, or chromatin modifying complexes such as Polycomb Repressor Complexes (PRC) 1 or 2, can alter the balance of HSC self-renewal and differentiation without causing HSC/MPPs to convert to a relatively homogenous lymphoid primed state.^42–47^ In contrast, simultaneous inactivation of MLL3 and MLL4 results in complete loss of myeloid-, erythroid- and megakaryocyte-biased progenitors, loss of self-renewal capacity and rapid conversion of essentially all Kit^+^ progenitors into a non-self-renewing B-cell-like population that is transcriptionally and functionally distinct from any normal B-cell progenitor. At a molecular level, MLL3/4 directly and redundantly regulate several transcription factors and other epigenetic regulators that are critical for HSC maintenance, and they thus sustain much of the HSC/MPP enhancer and superenhancer network.

Our results raise the question of why HSCs, MPPs and more committed progenitors all default to a B-cell-like state, uniformly, when MLL3 and MLL4 are inactivated. One possibility is that MLL3/4 target genes, such as *Runx1*, *Gata2* and *Mecom*, directly or indirectly antagonize B-cell master regulators such as *Pax5* and *Ebf1*. Consistent with this interpretation, *Gata2* has been shown to suppress expression of several B-cell-associated genes, including *Ebf1* and *Pax5*, in mice.^48^ Furthermore, deleting *Gata2b* in zebrafish leads to B-cell priming but impaired B-cell maturation.^49^ When *Mll3/4*-dependent enhancer and superenhancer networks collapse after conditional *Mll3/4* deletion, a feed forward circuit involving B-cell transcription factors (e.g., PAX5 and EBF1) could rapidly assert B-cell identity. Alternatively, MLL4 has been shown to partition chromatin and form phase-separated transcriptional condensates.^20^ EBF1 can also initiate phase-separated condensates.^50^ Loss of MLL3 and MLL4 could alter condensate formation to favor EBF1-mediated B-cell priming. In either scenario, HSC/MPP multipotency reflects constant tension between MLL3/4-dependent stemness programs and MLL3/4-independent B-cell programs.

Finally, this study reveals the degree to which MLL3 and MLL4 functions can be isolated from their histone methyltransferase activities. This observation aligns with prior work showing that MLL3 and MLL4 have SET-independent functions during early embryonic development,^25^ as well as studies showing that other chromatin modifying enzymes, including UTX, MLL1 and EZH2, can sustain normal hematopoiesis even when their catalytic domains are disabled.^51–53^ Of note, the gene expression changes observed in *Mll3/4* deficient progenitors are distinct from recently described changes in progenitors deficient for *Utx* and *Ptip*,^38,54–59^ both of which encode shared MLL3/4 COMPASS binding partners. Thus, MLL3 and MLL4 regulate gene expression independently of these binding partners, and vice versa.

## Acknowledgements

This work was supported by grants to J.A.M. from NHLBI (R01HL152180), NCI (R01CA285272), Gabriel’s Angel Foundation and the Mark Foundation, the Edward P. Evans Foundation and the Children’s Discovery Institute of Washington University and St. Louis Children’s Hospital, to G.A.C from NIDDK (R01DK124883) and to J.J.B. from NIAID (R01AI173077). H.C.W. is supported by the American Society of Hematology Graduate Award. J.A.M. is a Scholar of Blood Cancer United.

## Authorship

J.A.M. designed and oversaw all experiments, conducted experiments, interpreted data, wrote the manuscript with H.C.W., and secured funding. H.C.W and R.C. designed, conducted, and interpreted experiments. W.Y. and R. Z. performed bioinformatic analyses. R.M., T.H., R.M.P., E.B.C. and E.D. performed experiments and interpreted data. G.X. and K.G. provided critical reagents. G.A.C oversaw epigenomic experiments. J.J.B oversaw experiments related to B-cell development. All authors reviewed and edited the manuscript. Correspondence and requests for materials should be addressed to.

