## Supplemental Materials for "MLL3 and MLL4 sustain hematopoietic stem cell multipotency by opposing a B-cell default state"

### SUPPLEMENTAL INFORMATION

#### Supplemental Methods

##### Mouse strains and treatments

*Mll4*$^{f}$, *Mx1-Cre*, *Ubc-Cre::ER*, *LysM-Cre*, and *MBI-Cre* mouse lines were obtained from the Jackson Laboratory. *Mll3*$^{f}$, *Mll3*$^{Y4792A}$, and *Mll4*$^{Y5477A}$ mouse lines were previously described.[1,2] Male and female mice were used in similar numbers. Mice were housed in a standard pathogen free barrier facility, and all procedures were performed according to an IACUC approved protocol at Washington University School of Medicine. Intraperitoneal injections of pIpC (GE Life Sciences) were administered at 10 µg/day every other day for a total of 3 doses. Oral gavage of tamoxifen was administered at 100 mg/kg body weight every day for a total of 5 doses. For transplantation experiments, mice were gamma irradiated with two doses of 550 cGy spaced by 3 hours. Cells were then injected to the retroorbital sinus.

##### Flow cytometry and histology

Cells were isolated, stained, and analyzed as previously described.[1,3] Bone marrow cells were obtained by flushing the long bones (tibias and femurs) or by crushing long bones, pelvic bones and vertebrae with a mortar and pestle in calcium and magnesium-free Hank's buffered salt solution (HBSS), supplemented with 2% heat-inactivated bovine serum (Gibco). The cells were then stained sequentially, for 20 minutes each, with biotin conjugated anti-CD117 (c-kit; 2B8) and then Streptavidin-conjugated paramagnetic beads (Biolegend). CD117$^{+}$ cells were enriched by magnetic selection with LS magnetic columns (Miltenyi Biotec). Cells were stained for flow cytometry using the following antibodies, all of which were from Biolegend except as indicated: CD150 (TC15-12F12.2), CD48 (HM48-1), Sca1 (D7), c-kit (2B8), Ter119 (Ter-119), CD3 (17A2), CD11b (M1/70), Gr1 (RB6-8C5), B220 (RA3-6B2), CD8a (53-6.7), CD2 (RM2-5), CD45.1 (A20), CD45.2 (104), surface IgM (RMM-1), CD105 (MJ7/18), CD16/32 (93), CD71 (RI7217), CD43

(S11), CD19 (eBioscience, 1D3), IgD (11-26c.2a), CD21/35 (7E9), and CD23 (B3B4). Lineage stains included CD2, CD3, CD8a, Ter119, B220 and Gr1, or without B220 for B-cell analyses. The following surface marker phenotypes were used to define cell populations at all ages: HSCs ($CD150^+$, $CD48^-$, $Lineage^-$, $Sca1^+$, $c\text{-}kit^+$), MPPs ($CD48^+$, $Lineage^-$, $Sca1^+$, $c\text{-}kit^+$), pGM ($Lineage^-$, $Sca1^+$, $c\text{-}kit^+$, $CD150^-$, $CD105^-$, $CD16/32^-$), GMP ($Lineage^-$, $Sca1^+$, $c\text{-}kit^+$, $CD150^-$, $CD105^-$, $CD16/32^+$), pro-B ($Lineage^-$, $B220^{mid}$, $IgM^{lo}$, $CD43^+$), pre-B ($Lineage^-$, $B220^{mid}$, $IgM^{lo}$, $CD43^-$), recirculating B cells ($Lineage^-$, $B220^{hi}$), follicular B cells ($CD19^+$, $CD23^+$, $CD21/35^+$), marginal zone B cells ($CD19^+$, $CD23^-$, $CD21/35^+$), and transitional B cells ($CD19^+$, $CD23^-$, $CD21/35^-$, $IgD^-$, $IgM^+$). Non-viable cells were excluded from analyses by 4',6-diamidino-2-phenylindone (DAPI) staining (1 mg/ml). Flow cytometry was performed on a BD FACSAria Fusion flow cytometer (BD Biosciences) and analyzed by FlowJo. Cells were double sorted to ensure purity.

For histology studies, decalcified femurs were paraffin embedded, cut, and H&E stained according to standard protocols by the Musculoskeletal Histology and Morphometry Core at Washington University School of Medicine. For cytospin morphology assays, sorted $CD11b^+Gr1^+$ bone marrow cells were cytospun and Wright-Giemsa stained.

**Colony formation and B-cell potential assays**

To measure myeloid colony forming potential, single HSCs were sorted individually into wells of a 96-well plate and cultured in Methocult M3434 media (Stem Cell Technologies). Colonies were scored and harvested for genotyping 14 days later.

To measure B-cell potential, OP9 cells were seeded in 96-well plates at 12,500 cells per well in minimum essential medium, 20% FBS, and 1% penicillin-streptomycin. Limiting dilutions of LSK cells were then sorted into at least 24 wells per cell dose, with 3 biological replicates per genotype. Cells were cultured with FLT3L and IL-7 (10 ng/ml), and media was changed every 4 days. After 12-14 days, each well was analyzed by flow cytometry for the presence of $CD19^+$ or

B220[+] B-cells. Wells were considered positive for B-cell potential if B-cells were detected after the culture period. Extreme Limiting Dilution Analysis (ELDA)[4] was used to compare limiting dilution curves.

**32D cell culture and CRISPR editing**

32D cells were cultured in RPMI 1640 medium with 10% FBS, 1% penicillin-streptomycin, and 10ng/ml IL-3, at 37°C and 5% $CO_2$.

To introduce a 3xFLAG epitope after the initiator methionine at the endogenous *Mll3* locus, ribonucleoprotein complex (RNP) was prepared by incubating Cas9 (IDT) with the gRNA targeting the first exon of *Mll3* (5'- CCUGGUGACUAGGAUGUCGU-3') and electroporated into 32D cells with an ssODN that carried the 3xFLAG sequence (5'-GGCGGCTGCTGCTGCTCCGCGCTCCGGTCCTCCTCCGACGACTTGTCATCGTCGTCTTTG TAGTCGATGTCGTGATCCTTATAATCGCCGTCGTGGTCCTTGTAGTCCATCCTAGTCACCAG GAAAGACACATGGATCCCCGTCCTC-3'). Fifty thousand 32D cells were resuspended in buffer R (Thermo) and electroporated with 1600V, 20ms, 1 pulse using a Neon Transfection kit (Thermo). Single cells were sorted on 96-well plates and screened by PCR to identify insertions. Clones with homozygous FLAG insertions were confirmed by Sanger sequencing. Primers used to identify successfully targeted clones were 5'- gacggggatccatgtgtcttt-3' and 5'-ccctcgcgctcaacttactt-3'.

To generate *Mll3*$^{-/-}$ cells from the 3xFLAG-MLL3 32D line, RNP was prepared by mixing Cas9 with gRNAs targeting exon 14 of *Mll3* (5'-GUCACUCCAAAAAUUGGCAU-3'). Isogenic 3xFLAG-MLL3 32D cells were electroporated as described above. Single cells were sorted on 96-well plates and screened by next-generation sequencing. The primers used to identify clones with null mutations were 5'- gcaactgacctgaacaaggc-3' and 5'-acatcaccctacctgtttgga-3'.

**Co-Immunoprecipitation and Western blot**

For co-Immunoprecipitation with the FLAG epitope, 32D cells were washed with cold PBS and incubated on ice for 15 minutes in cold Buffer A (10mM HEPES, 10mM KCl, 0.1mM EDTA, 0.1mM EGTA, and protease inhibitor). 10% IGEPAL was added, and lysates were centrifuged at 14,000rpm for 5 minutes at 4°C. Nuclei were washed with cold PBS and resuspended in modified RIPA buffer (0.1% IGEPAL, 0.1% sodium deoxycholate, 150mM NaCl, 50mM Tris pH 8, 1mM $MgCl_2$, 1U/µl Benzonase, and protease inhibitor). Nuclei were incubated on ice for 30 minutes and debris was removed by centrifugation at 14,000rpm for 15 minutes. Protein concentrations were measured by the BCA assay (Thermo Fisher Scientific). Nuclear lysates were incubated with FLAG (M2) magnetic beads (Sigma-Aldrich) overnight at 4°C with rotation. Beads were washed twice with cold wash buffer (150mM NaCl, 50mM Tris pH 8, and protease inhibitor) with 1% IGEPAL and then twice without IGEPAL. Immunoprecipitated proteins were eluted using 3xFLAG peptide (Sigma-Aldrich) and analyzed by Western Blotting as previously described.[1]

For analyses of histone modifications, COMPASS subunits, and surface IgM, 30,000 $Lineage^-Kit^+Sca\text{-}1^+$ cells were sorted directly into 10% Trichloroacetic acid. The precipitates were pelleted, washed with acetone, and solubilized before running on NuPAGE 4-12% Bis-Tris gels or 3-8% Tris-Acetate gels (Thermo). Proteins were transferred to PVDF membranes and Western Blotting was performed as previously described.[1]

**CITE-seq library construction, sequencing, and analysis**

We isolated $CD45.2^+Lineage^-Kit^+Sca\text{-}1^+$ (LSK) cells by flow cytometry and stained with Total-Seq B antibodies (Biolegend) to CD117, CD150, CD48, CD135, CD34, and CD201. The experiments using $CD45.2^+Kit^+$ cells were performed with two unpooled biological replicates each stained with a unique Total-Seq B Hashtag antibody. All $Kit^+$ cells were stained with Total-Seq B antibodies (Biolegend) to CD117, Sca-1, CD150, CD48, CD135, CD16/32, CD127, CD41, B220, Gr1, and CD11b. Libraries were generated with 10x Genomics Chromium Next GEM Single Cell 3′ v3.1, Chromium Single Cell 3’ Feature Barcode Library, Dual Index NT Set A, and Dual Index

TT Set A kits according to the manufacturer's instructions. Libraries were sequenced on an Illumina Novaseq X Plus. The Cell Ranger v.6.0.1 pipeline (10x Genomics) was used to process data. Digital gene expression (DGE) files were then filtered and normalized as previously described.[3] The AltAnalyze toolkit was then used to perform Iterative Clustering and Guide-gene Selection (ICGS, version 2).[5,6] Cluster identities were annotated based on marker gene expression, as defined by ICGS, and by immunophenotypes based on antibody directed tags (ADT). Thresholds for ADT positive and negative expression are shown in Supplemental Figures S3-4.

Differential expression analyses were performed using pseudoreplicates (n=2 per biological replicate, n=2 biological replicates per genotype, n=4 total) that were generated using Presto (https://github.com/immunogenomics/presto). Differentially expressed genes were then identified among the pseudoreplicates using DESeq2 and Gene Set Enrichment Analysis (GSEA).[7,8] Quadratic programming was performed using approaches developed by Morris and colleagues.[9] We first identified genes that distinguished the most immature HSC/MPP cluster (cluster 28 in Figure 2G) from the most differentiated myeloid clusters (clusters 22 and 30 in Figure 2G), based on pseudobulk gene expression comparisons. Once the list of differentially expressed genes was generated, we used Capybara to perform quadratic programming and assign myeloid identity scores to all cells in the annotated HSC/MPP, GMP (cluster 11) and myeloid clusters. Gene Set Variation Analysis (GSVA) was performed with the curated B-cell signature shown in Supplemental Table S3 using a Gaussian kernel and normalized gene expression from ICGS.[10] For Figure 7, we used Symphony[11] to assign cells to clusters using the same cluster identities annotated in Figure 2G.

**5' scRNA-seq library construction, sequencing, and analysis**

We isolated CD45.2$^+$B220$^{lo}$ and CD45.2$^+$Kit$^+$ cells by flow cytometry and generated libraries with 10x Genomics Chromium Next GEM Single Cell Single Cell 5′ v2 according to the manufacturer's instructions. Libraries were sequenced on an Illumina Novaseq X Plus. The Cell Ranger v7.1.0 pipeline was used to align, filter and normalize digital gene expression files. The AltAnalyze toolkit was then used to perform Iterative Clustering and Guide-gene Selection (ICGS, version 2) and annotate the identified clusters to cell types. The "cellranger multi" output in "filtered_contig_annotations.csv" was parsed to get the IgH/L identifications. Seurat was used to estimate the cell cycle scores and assign cell cycle status.[12]

**Single-cell ATAC-seq library construction, sequencing, and analysis**

We sorted CD45.2$^{+}$Lineage$^{-}$Sca1$^{+}$Kit$^{+}$ cells by flow cytometry, isolated nuclei and generated libraries with 10x Genomics Chromium Next GEM Single Cell ATAC Kits v2 according to the manufacturer's instructions. Libraries were sequenced on an Illumina Novaseq X Plus. The Cell Ranger ATAC v2.1.0 pipeline was used to align raw sequencing reads and generate BAM and fragment files. ArchR[13] was used to filter doublets, cluster cells and call peaks. Gene expression data from CITE-seq were integrated based on concordance between gene accessibility (Gene Score Matrix) and gene expression (Gene Integration Matrix). Supercluster identities from Figure 2A were used to annotate ArchR clusters. Cluster-specific elements were defined as marker peaks in each supercluster using ArchR and were aggregated by genotype to generate histograms as shown in Figures 4E, 5A and S7D. Cluster-specific elements were aligned with previously described H3K4me1 and H3K27ac ChIPmentation data from purified HSC and MPP cells (GSE158159), as well as LSK and GMP cells (GSE243869).[1,3] Motif enrichment in differentially accessible regions was calculated by ArchR based on the cisbp motif set and also by ChromVAR.[14] Motif footprinting and Peak2Gene (P2G) linkage analyses were also performed using the respective ArchR functions. In panels showing aggregate scATAC-seq peaks, reads were exported from ArchR and aggregated into pseudoreplicates, then mapped to WashU Epigenome browser.

**ChIPmentation and ChIP-seq analyses**

For H3K27ac ChIPmentation, 100,000 LSK cells were double sorted into PBS with 0.1% BSA, crosslinked with 1% formaldehyde, and quenched with 125mM glycine. For FLAG ChIPmentation, 20 million 32D cells were crosslinked with 1% formaldehyde and 2mM disuccinimidyl glutarate (DSG) for 10 minutes at room temperature, then quenched with 125mM glycine. Cells were sonicated using a Covaris E220 to achieve fragment sizes around 200-500 bp. Sheared chromatin was incubated with H3K27ac antibody (Diagenode) or FLAG (M2) antibody (Sigma-Aldrich) overnight. Libraries were prepared using the ChIPmentation protocol as previously described,[3] and sequenced on a NovaSeq X Plus. Reads were mapped to mm10, and peaks were called with MACS2. Signal tracks were visualized with the WashU Epigenome browser.[15] HOMER was used to identify transcription factor binding sites that were enriched within MLL3-bound regions.[16]

Analysis of superenhancers was performed with Rank Ordering of Super Enhancers (ROSE).[17] The DiffBind R package was used to perform differential binding analysis across sample groups. ChIP-seq data in pro-B cells on EBF1 (GSE158673), PAX5 (GSE140975), and MED1 for superenhancers (GSE44288) were previously described.[17-19] H3K4me1 and H3K27ac ChIPmentation data for control or *Mll3*$^{\Delta/\Delta}$ HSC, MPP, LSK and GMP have been previously described (GSE158159, GSE243869).[1,3]

**CUT&RUN library construction, sequencing, and analysis**

CUT&RUN was performed using the EpiCypher CUTANA CUT&RUN Kit (Cat. #14-1048, version 5) according to manufacturer instructions. 500,000 32D cells were lightly fixed with 0.1% formaldehyde for 1 minute and isolated nuclei were immobilized using concanavalin A beads. Cells were incubated with homemade MLL3 or MLL4 antibodies overnight (Kai Ge lab), followed by targeted chromatin digestion and release by pAG-MNase. The CUT&RUN enriched DNA was purified and used for library preparation using the CUTANA CUT&RUN Library Prep Kit per manufacturer instructions (Cat. #14-1001). The end repair reaction was performed on the purified DNA, followed by adaptor ligation, uracil excision and library amplification. DNA was purified and sequenced on a NovaSeq X Plus targeting 8 million pair-end reads per sample.

Sequencing reads were processed with CUT&RUNTools 2.0[20] using default parameters, except that BAM files were not filtered by the 120-bp fragment size criterion. The pipeline included read preprocessing, genome alignment, normalization, and peak calling after subtracting IgG signals as background. Peaks identified by MACS2, together with the corresponding signal tracks, were used for downstream interpretation and analysis. Each group contained 4 technical replicates. Reads were processed per replicate to obtain individual alignments, then pooled to generate signal tracks and peaks for each group. To identify common peaks among the MLL3/4 CUT&RUN and FLAG-MLL3 ChIPmentation, the Venn diagram was generated by pooling peaks from all 3 datasets to get the combined regions first, before counting overlapping with the 3 individual sets of peaks.

**Statistical analysis**

Group sizes are indicated in the figure legends. Groups were compared with one way ANOVA and Holm-Sidak posthoc test to correct for multiple comparisons. CITE-seq, scATAC-seq, and 5'RNA-seq analyses are described above.

**Data availability**

Datasets generated during this study are available at Gene Expression Omnibus under accession numbers GSE301672, GSE301674, GSE301675, GSE301676, GSE301677, GSE301678, GSE301679, and GSE301680.

#### Supplemental Figures

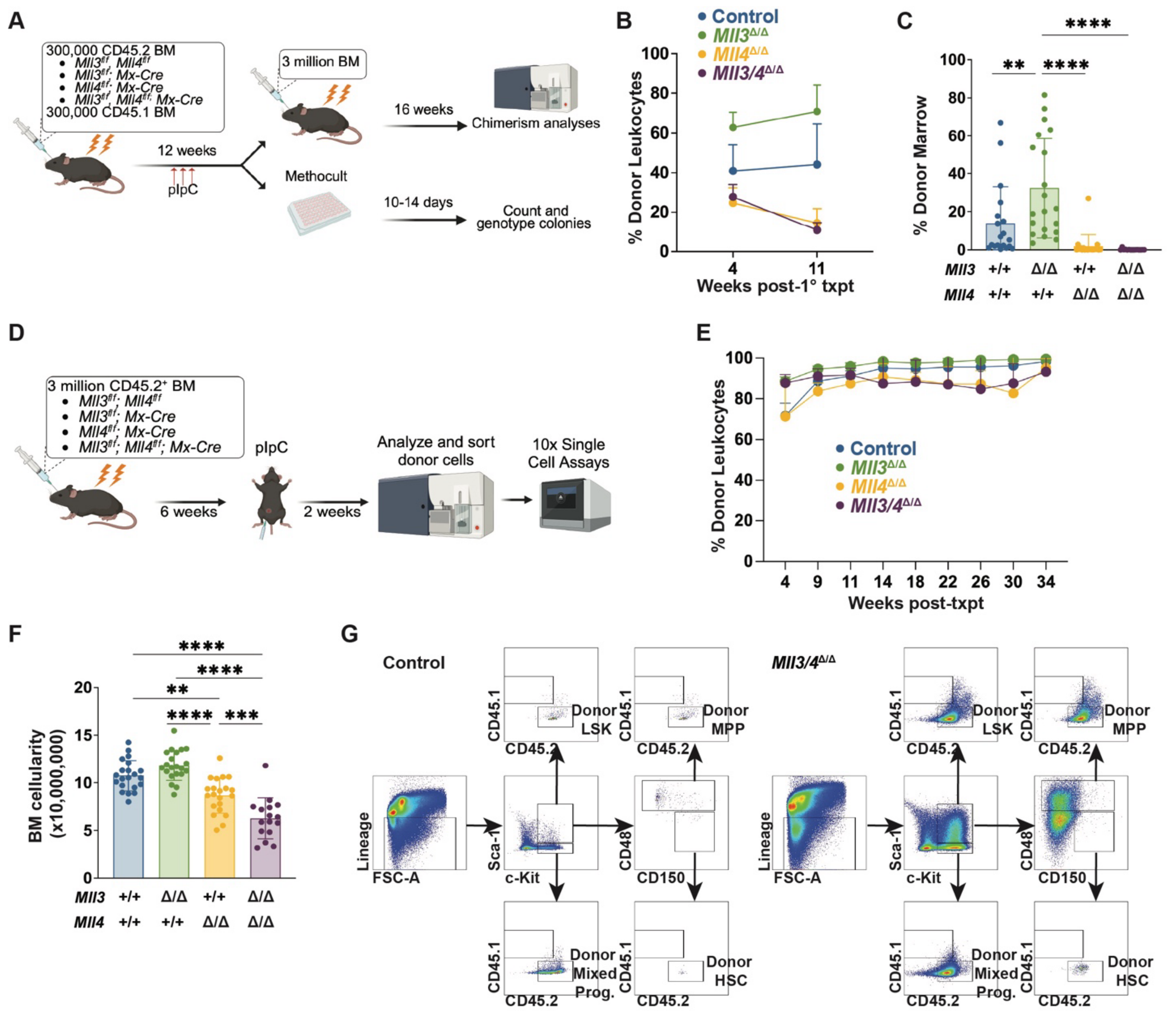


**Figure S1. Competitive and non-competitive transplantations of *Mll3/4* conditional knockout bone marrow, related to Figure 1.** (A) Overview of serial transplantation. Equal numbers of CD45.2$^+$ bone marrow cells of the indicated genotypes and CD45.1$^+$ competitor cells were transplanted into lethally irradiated recipients, and 3 doses of pIpC were administered 6-week post-transplantation. Three million bone marrow were harvested at 6 weeks post-pIpC and transplanted into lethally irradiated secondary recipients to assess repopulating activity. At the time of the primary transplant takedown, single HSCs were sorted per well on 96-well plates with Methocult M3434 for genotyping. (B) Peripheral blood chimerism in primary transplantation recipients of the indicated genotypes, prior to pIpC treatment. (n=10-15). (C) Bone marrow chimerism in secondary transplantation recipients of the indicated genotypes at 16 weeks post-transplant. (n=15-20). (D) Overview of noncompetitive transplantation strategy to assess population sizes and obtain cells for CITE-seq/scATAC-seq. Flow cytometry and single cell analyses were performed at 2 weeks after pIpC treatment. A second cohort of recipients was observed until *Mll3/4*$^{\Delta/\Delta}$ mice became moribund around 29-30 weeks after pIpC treatment. At that point, the mice were euthanized along with littermate controls and single mutant recipients for bone marrow evaluation and histology. (E) Peripheral blood donor (CD45.2) chimerism in recipients of the indicated genotypes. (n=10-15). (F) Bone marrow cellularity in recipients of the indicated genotypes at 2 weeks after pIpC treatment. (n=16-21). (G) Gating strategy for donor HSC, MPP, and Lineage$^{-}$Kit$^{+}$Sca1$^{-}$ progenitor populations in control and *Mll3/4*$^{\Delta/\Delta}$ recipients. For all panels, error bars reflect standard deviation, *p<0.05, **p<0.01, ***p<0.001 as calculated by one-way ANOVA followed by Holm-Sidak posthoc test.

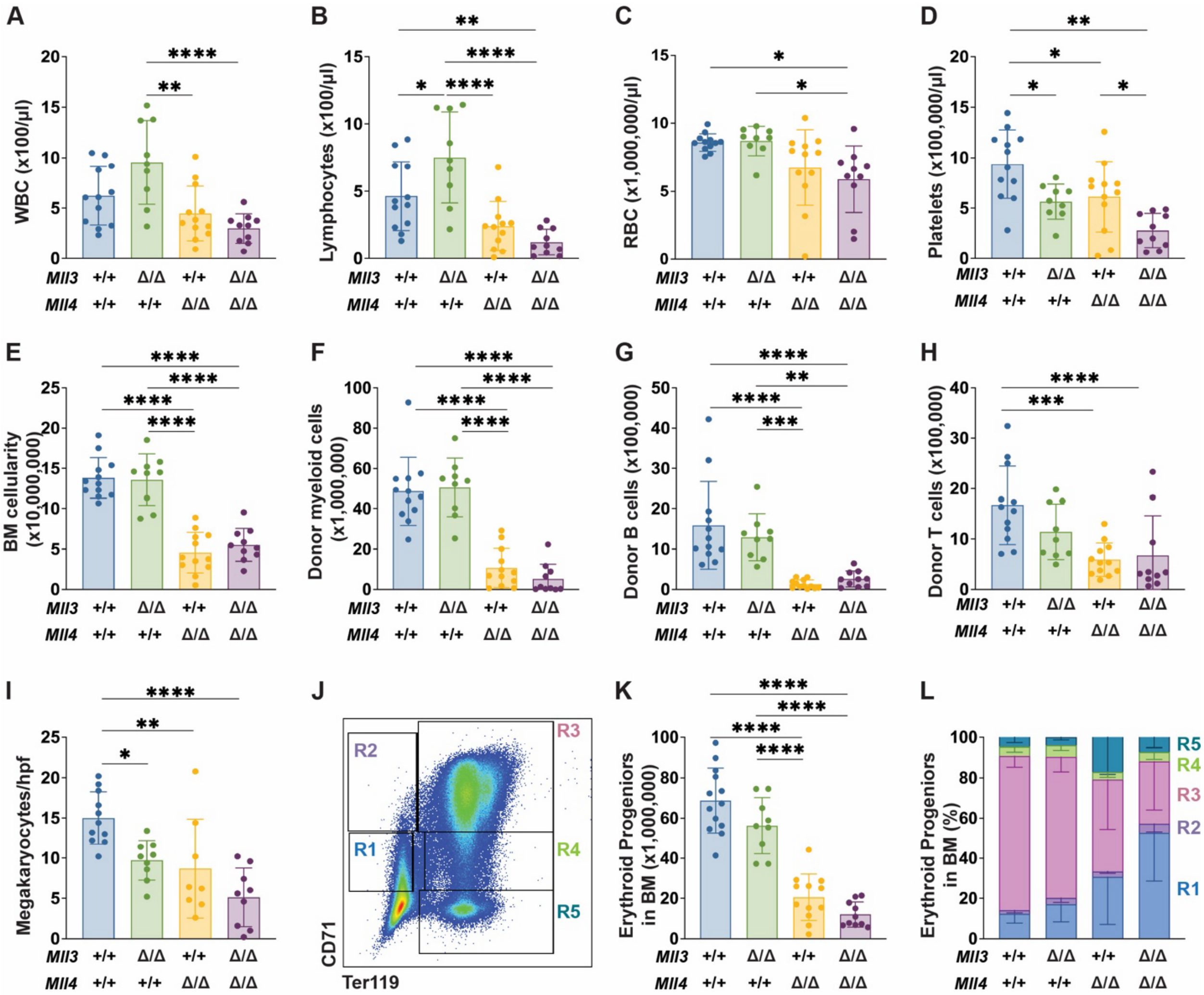

**Figure S2. *Mll3* and *Mll4* deletions result in bone marrow failure 30 weeks after pIpC treatment, related to Figure 1.** (A-D) White blood cell (WBC), lymphocyte, red blood cell (RBC), and platelet counts in peripheral blood of recipients of the indicated genotypes. (n=9-12). (E-H) Total bone marrow cell counts, donor (CD45.2) myeloid, B cell, and T cell numbers in the bone marrow of recipients of the indicated genotypes. (n=9-12). (I) Megakaryocyte counts per high power field of H&E bone marrow sections. (n=8-11). (J) Gating strategy for bone marrow erythroid progenitors. R1 represents proerythroblasts, R2 early basophilic erythroblasts, R3 late basophilic erythroblasts, R4 orthochromatophilic erythroblasts, and R5 reticulocytes. (K, L) Total erythroid progenitor numbers and distribution of erythroid progenitor populations in the bone marrow of recipients with the indicated genotypes. (n=9-12). For all panels, error bars reflect standard deviation, *p<0.05, **p<0.01, ***p<0.001 as calculated by one-way ANOVA followed by Holm-Sidak posthoc test.

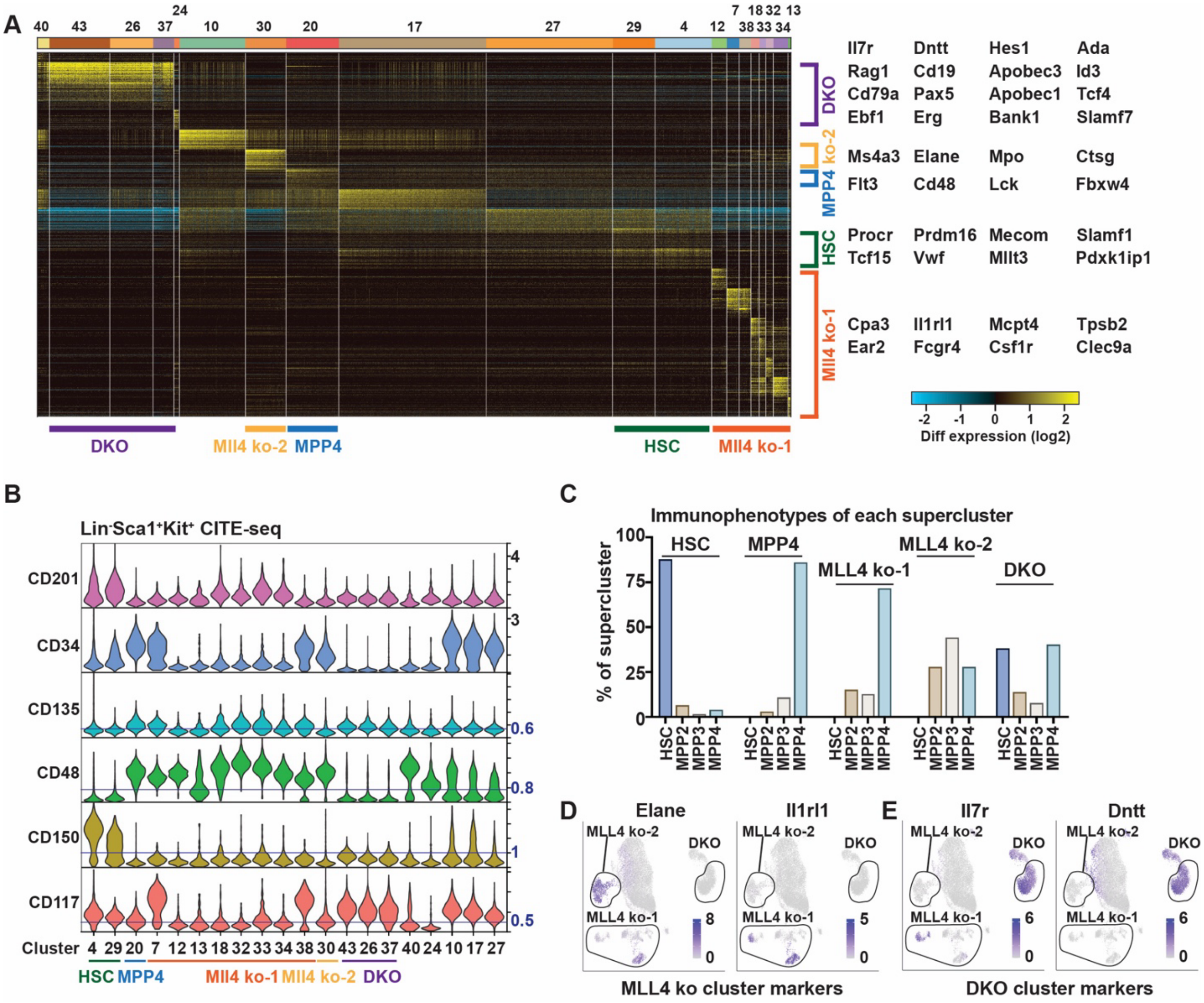


**Figure S3. Compound *Mll3* and *Mll4* deletions cause hematopoietic progenitors to convert to a B-cell-like default state, related to Figure 2.** (A) Heatmap and clustering of LSK cells based on CITE-seq gene expression. Clustering was performed with ICGS, version 2. A subset of marker genes used to define cluster identities are shown to the right. Superclusters are indicated, in some cases by joining clusters from adjacent clades. (B) Violin plot showing antibody directed tag (ADT) signal in each cluster for the LSK CITE-seq experiment. Thresholds defining positive and negative populations are indicated for markers that were used to define immunophenotypic populations. Supercluster identities are annotated below the cluster numbers. (C) HSC, MPP2, MPP3 and MPP4 immunophenotype frequencies in each supercluster. (D, E) Expression of myeloid and lymphoid marker genes.

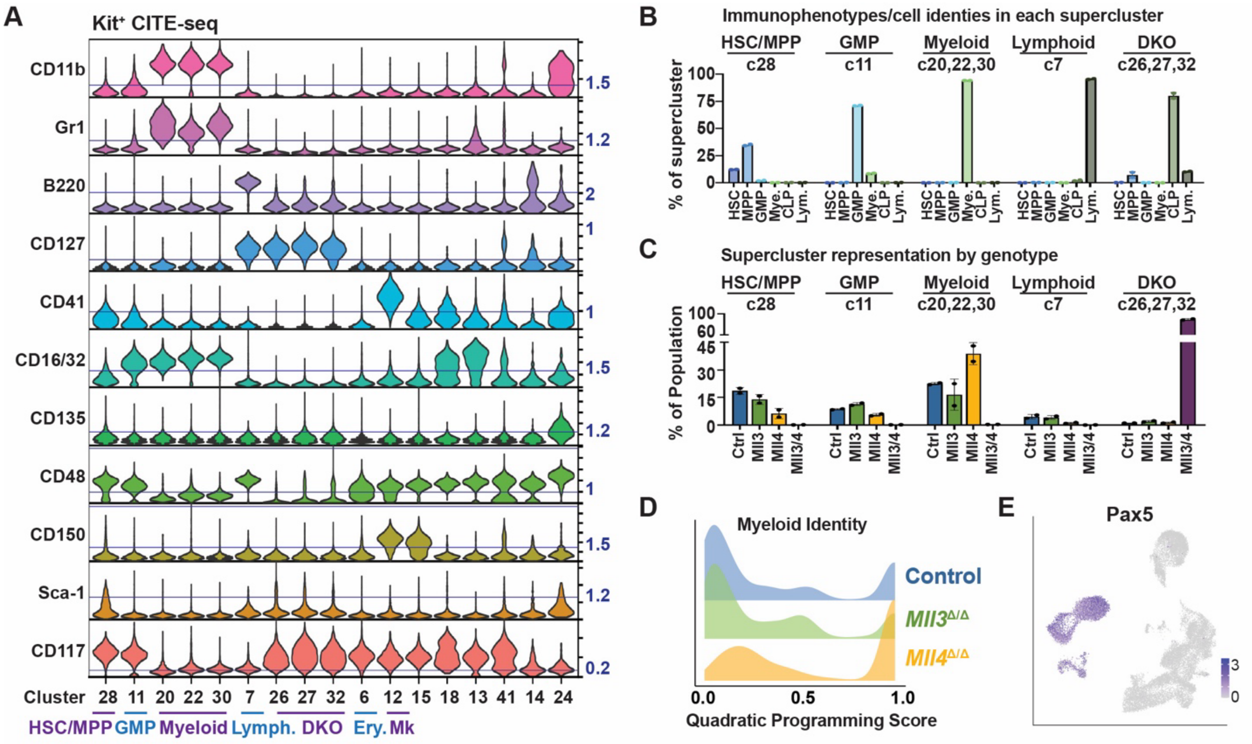


**Figure S4. Compound *Mll3* and *Mll4* deletions cause hematopoietic progenitors to convert to a B-cell-like default state, related to Fig. 2.** (A) Violin plot showing antibody directed tag (ADT) signal in each cluster for the Kit$^+$ CITE-seq experiment. Thresholds defining positive and negative populations are indicated for markers that were used to define immunophenotypic populations. Supercluster identities are annotated below the cluster numbers. (B) HSC, MPP, GMP, myeloid, CLP, or lymphoid immunophenotype frequencies in each supercluster. (C) Supercluster distributions for indicated genotypes. (D) Distribution of myeloid identity quadratic programming scores for all HSC/MPP, GMP and myeloid supercluster cells combined for the indicated genotypes. (E) Expression of *Pax5* in individual cells.

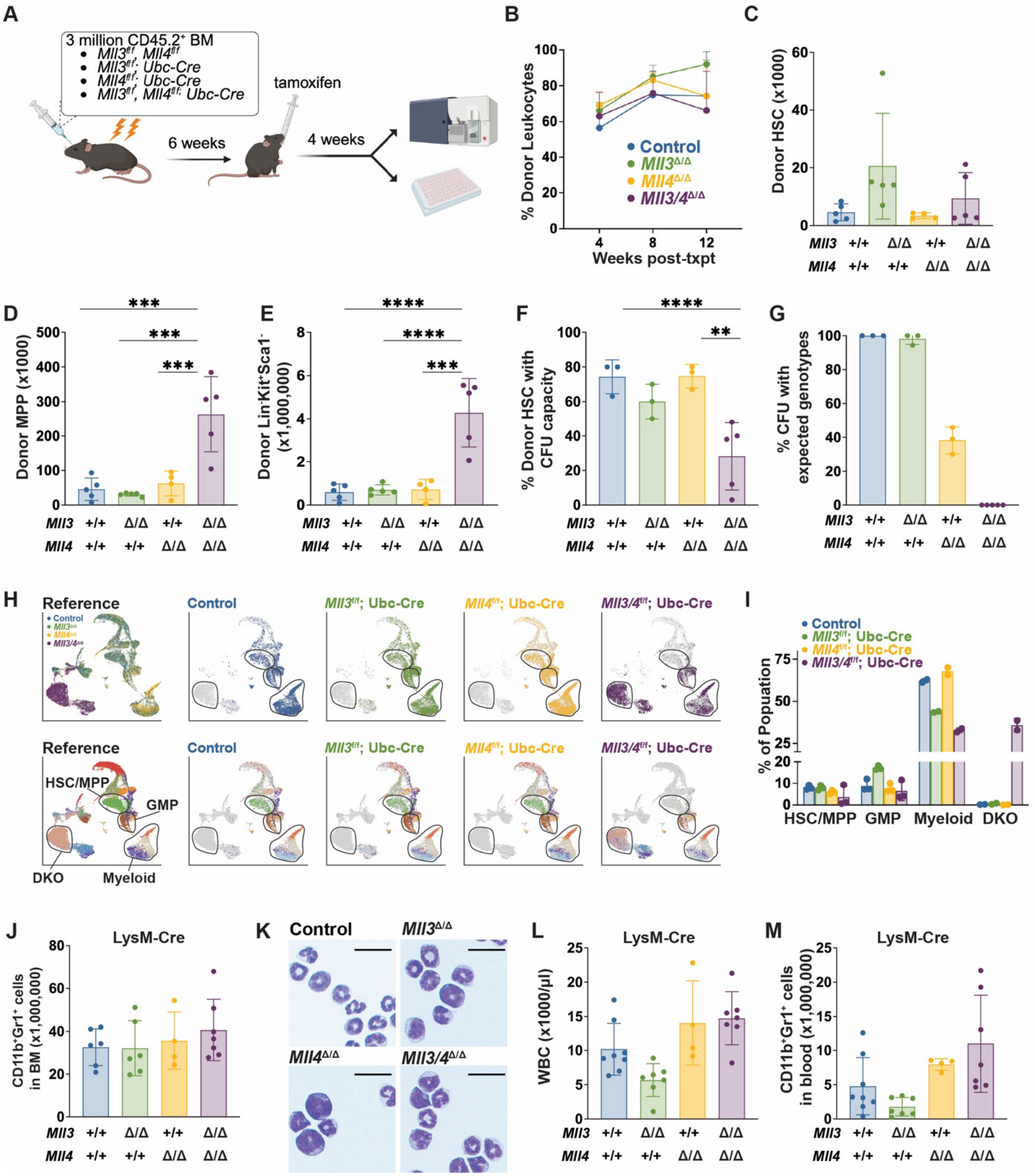


**Figure S5. MLL3 and MLL4 are necessary for myeloid commitment but dispensable in late myeloid maturation, related to Fig. 2.** (A) Overview of Ubc-CreER noncompetitive transplantation. Donor cells were analyzed by flow cytometry, and single donor HSCs per well were plated in a 96-well plate with MethoCult M3434 for genotyping. (B) Peripheral blood chimerism in recipients of the indicated genotypes. (n=4-5). (C-E) Donor HSC, MPP, and Lineage$^-$Kit$^+$Sca1$^-$ numbers in recipient bone marrow 4 weeks after tamoxifen treatment. (n=4-5). (F) Percentage of donor HSCs with myeloid colony-forming potential and (G) percentage of myeloid colonies with genotypes that are expected following complete deletion of floxed alleles. (n=3-5). (H) Distribution of single cell transcriptomes of *Mll3/4*$^{\Delta/\Delta}$ *Ubc-CreER* cells projected against *Mll3/4*$^{\Delta/\Delta}$ *Mx-Cre* Kit$^+$ cells from Figure 2G by Symphony. Cells are colored by genotype (top) or reference cluster identity (bottom) as in Figure 2G. (I) Percentage of cells for each genotype that map to HSC/MPP, GMP, myeloid and DKO superclusters by Symphony. Each dot reflects a pseudoreplicate. (J, K) Number and morphology of CD11b$^+$Gr1$^+$ cells in *LysM-Cre* bone marrow. (n=4-7). Scale bars represent 50μm. (L) White blood cell (WBC) count and (M) CD11b$^+$Gr1$^+$ myeloid cells in the peripheral blood of *LysM-Cre* mice of the indicated genotypes. (n=4-7). Error bars reflect standard deviation, *p<0.05, **p<0.01, ***p<0.001, ****p<0.0001 by one-way ANOVA and Holm-Sidak posthoc test.

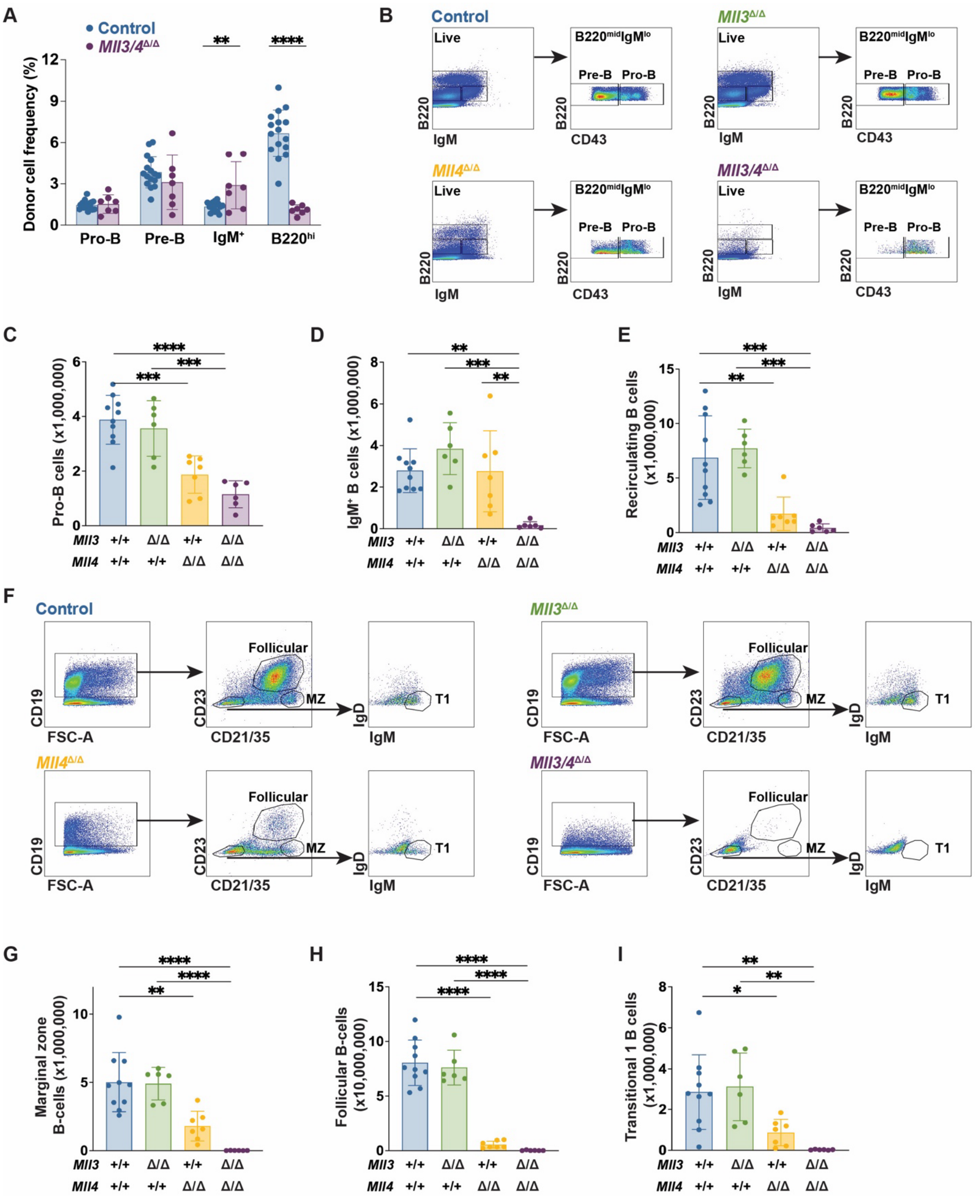


**Figure S6. MLL3 and MLL4 are required for B lymphopoiesis, related to Figure 3.** (A)

Frequency of donor pro-B, pre-B, IgM$^{+}$, and B220$^{hi}$ recirculating B cells in the bone marrow of control or *Ml3/4*$^{\Delta/\Delta}$ recipients from the noncompetitive transplantation illustrated in Figure S1H. (B) Representative flow cytometry plots and gating strategies for pro-B, pre-B, IgM$^{+}$, and B220$^{hi}$ recirculating B cells in the bone marrow of *MBI-Cre* conditional knockout mice. (C-E) Number of pro-B, IgM$^{+}$, and B220$^{hi}$ recirculating B cells in the bone marrow of *MBI-Cre* conditional knockout mice of the indicated genotypes. (n=6-10). (F) Representative flow cytometry plots and gating strategies for marginal zone (MZ), follicular, and T1 B cells in the spleen of *MBI-Cre* conditional knockout mice. (G-I) Numbers of marginal zone, follicular and transitional 1B-cells in the spleens of *MBI-Cre* conditional knockout mice of the indicated genotypes. (n=6-10). For all panels, error bars reflect standard deviation, *p<0.05, **p<0.01, ***p<0.001 as calculated by one-way ANOVA followed by Holm-Sidak posthoc test.

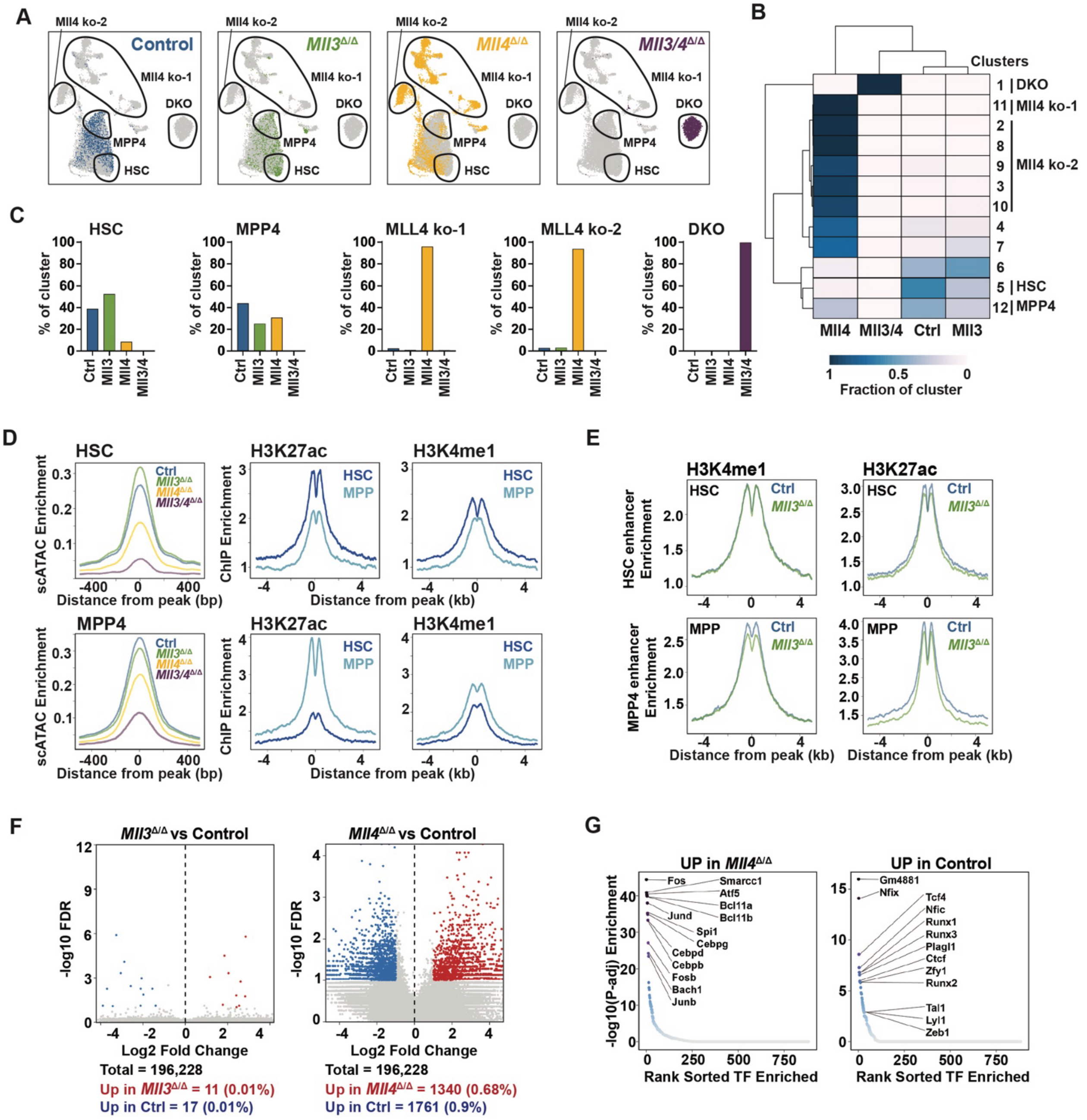


**Figure S7. scATAC-seq characterization, related to Figure 4.** (A) UMAPs representing scATAC-profiles of control, *Mll3*$^{\Delta/\Delta}$, *Mll4*$^{\Delta/\Delta}$, and *Ml3/4*$^{\Delta/\Delta}$ LSK. Superclusters were defined using integrated CITE-seq data from Figure 2A and are outlined in black. (B) Heatmap showing cluster and supercluster representation for each genotype. (C) Percentage of control, *Mll3*$^{\Delta/\Delta}$, *Mll4*$^{\Delta/\Delta}$, and *Ml3/4*$^{\Delta/\Delta}$ LSK in each scATAC-seq supercluster. (D) Left: Aggregate scATAC-seq HSC and

MPP4 supercluster-specific peaks by genotype. Middle and right: Aggregate H3K27ac and H3K4me1 signals from purified HSC or MPP cells at HSC and MPP4 supercluster-specific elements. (E) Aggregate H3K4me1 and H3K27ac levels in control and *Mll3*$^{\Delta/\Delta}$ HSCs and MPPs at HSC-specific (top) and MPP4-specific (bottom) cis-regulatory elements. (F) Volcano plots of differentially accessible regions between control and *Mll3*$^{\Delta/\Delta}$ cells (left), and between control and *Mll4*$^{\Delta/\Delta}$ cells (right). (G) Motif enrichment analysis on differentially accessible regions from control and *Mll4*$^{\Delta/\Delta}$ cells.

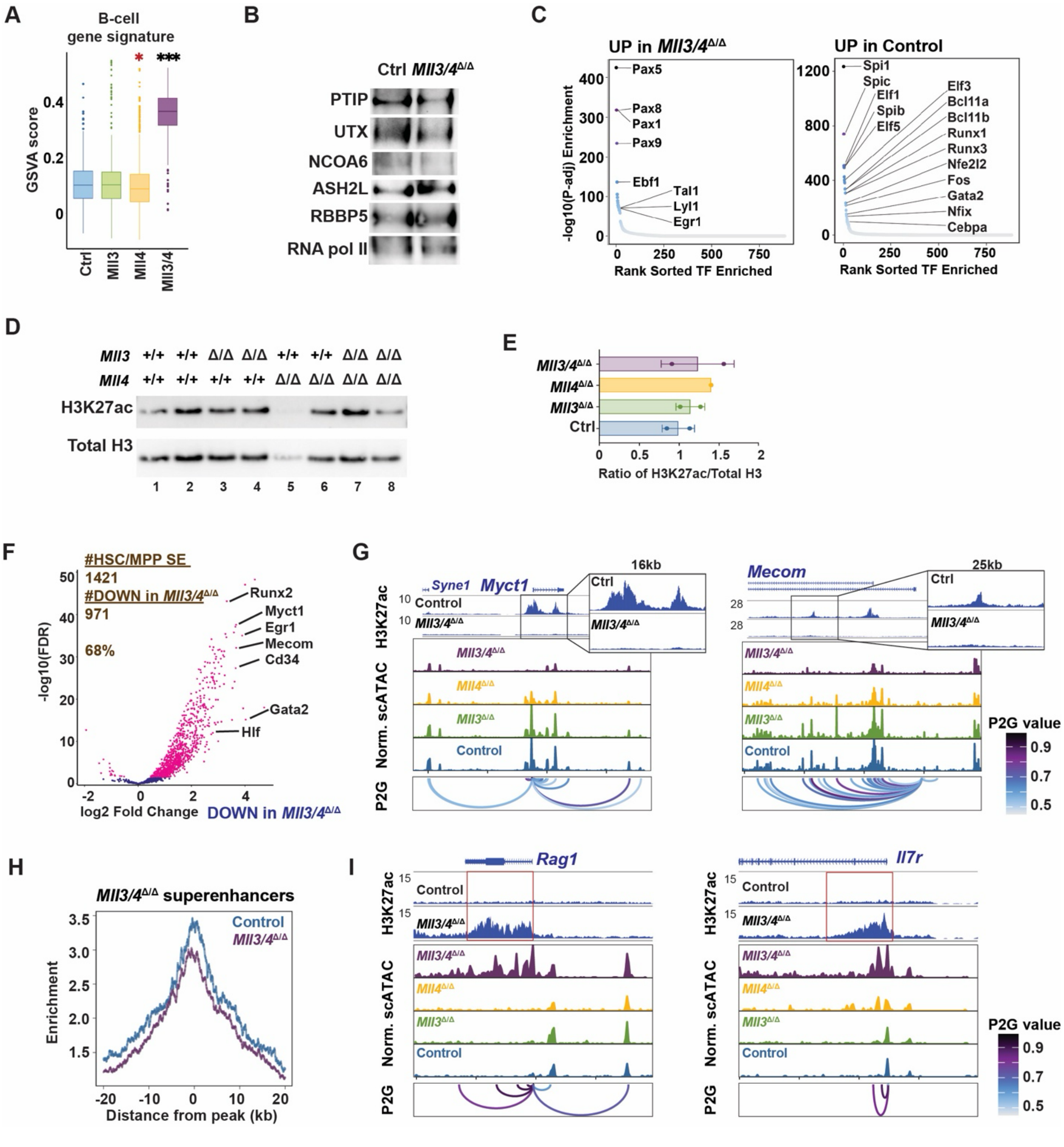


**Figure S8.** ***Mll3*** **and** ***Mll4*** **deletions cause HSC/MPP superenhancer inactivation and ectopic pro-B-cell superenhancer activation, related to Figure 5.** (A) B-cell signature enrichment in the DKO scATAC-seq cluster. *p<0.05, ***p<0.0001 by Wilcoxon test relative to control. (B) Western blot showing MLL3/4 COMPASS cofactor protein expression in control and *Ml3/4*$^{\Delta/\Delta}$ Lineage$^-$Sca1$^+$Kit$^+$ (LSK) cells, with RNA polymerase II (pol II) as control. (C) Motif enrichment in differentially accessible regions from *Mll3/4*$^{\Delta/\Delta}$ and control LSK. (D) Western blots showing H3K27ac and total Histone H3 in LSK cells of the indicated genotypes. Lane 5 had insufficient loading. (E) Quantification of the ratio of H3K27ac to total H3 band intensity from the Western blot in panel D, excluding lane 5. (n=1-2). There was no statistical difference between control and *Ml3/4*$^{\Delta/\Delta}$ H3K27ac/Total H3 ratios based on two-tailed Student's t-test. (F) Volcano plot demonstrating HSC/MPP superenhancers with significantly altered H3K27ac levels between control and *Mll3/4*$^{\Delta/\Delta}$ LSK cells. (G) Tracks showing reduced accessibility and H3K27ac at *Myct1* and *Mecom* in *Mll3/4*$^{\Delta/\Delta}$ LSK. Peaks to gene (P2G) indicates association between peaks and the promoter. (H) Aggregate H3K27ac levels at all *Mll3/4*$^{\Delta/\Delta}$ superenhancers. (I) Tracks showing increased accessibility and H3K27ac at *Rag1* and *Il7r* in *Mll3/4*$^{\Delta/\Delta}$ LSK. Red boxes indicate MLL3/4-independent superenhancers.

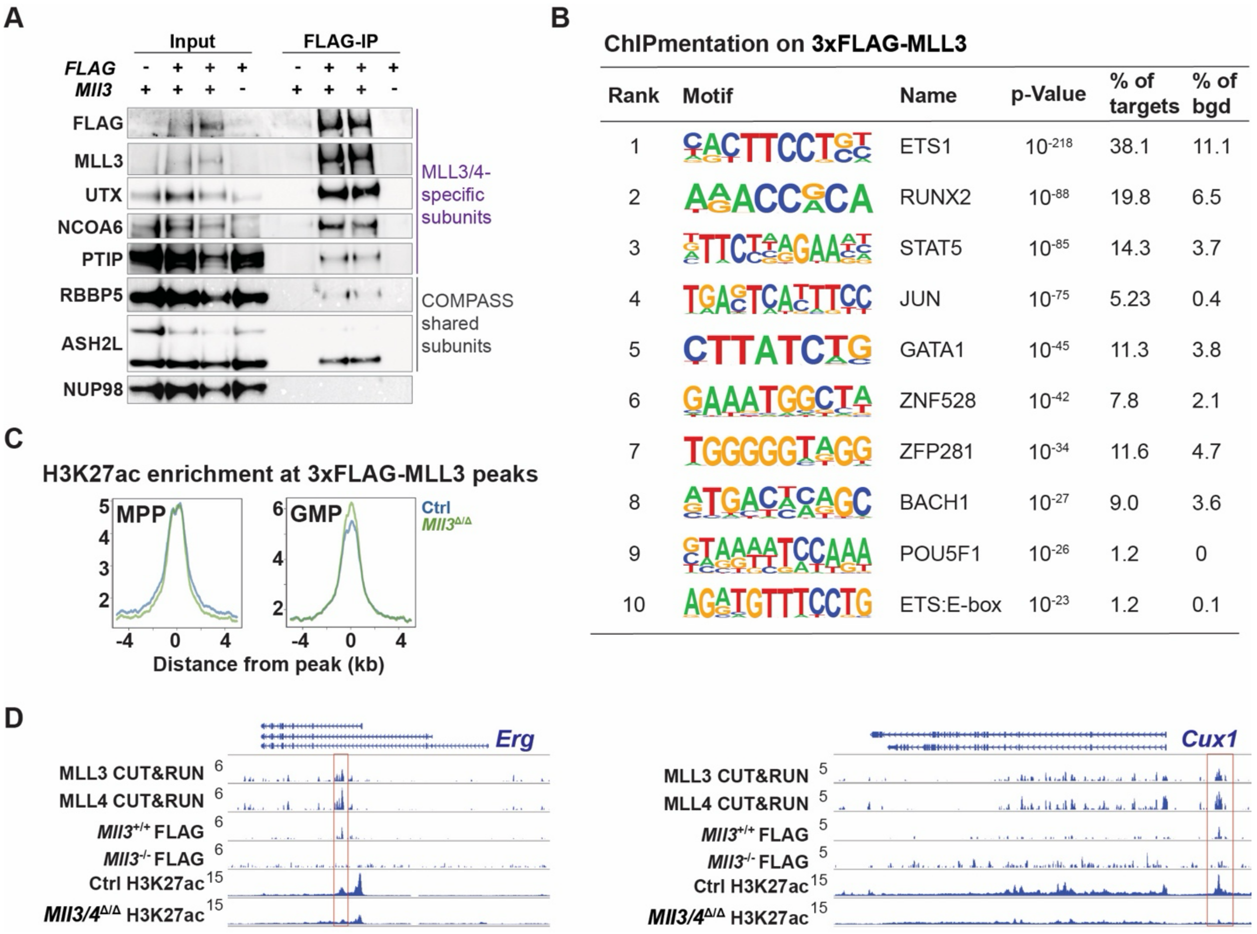


**Figure S9. MLL3 and MLL4 redundantly regulate a subset of HSC/MPP transcription factors, related to Figure 6.** (A) Western blot showing expression of COMPASS cofactors in inputs and eluates from 3xFLAG-MLL3 co-immunoprecipitation assays (FLAG antibody), with NUP98 as loading control. (B) Table of top 10 transcription factor binding motifs enriched in MLL3- and MLL4-bound regions, based on HOMER analysis. (C) Aggregate H3K27ac signals from purified control or *Mll3*$^{\Delta/\Delta}$ MPP and GMP cells at MLL3- and MLL4-bound peaks. (D) Representative tracks showing MLL3/4 CUT&RUN, FLAG-ChIPmentation, and H3K27ac at the *Erg* and *Cux1* loci. Red boxes indicate MLL3/4-bound enhancers.

#### Supplemental Tables

**Supplemental Table S1:** Quality control parameters for single cell assays.

**Supplemental Table S2:** Differentially expressed genes in *Mll4*$^{\Delta/\Delta}$ and *Mll3/4*$^{\Delta/\Delta}$ LSK cells.

**Supplemental Table S3:** B cell signature genes for Gene Set Variant Analysis.

**Supplemental Table S4:** Cluster-specific cis-regulatory elements.

**Supplemental Table S5:** Motif enrichment in *Mll3* and *Mll4*-dependent regulatory elements.

**Supplemental Table S6:** HSC, MPP, and MLL3/4-independent superenhancers.

**Supplemental Table S7.** MLL3- and MLL4-bound regulatory elements.
